# Generation and attenuation of variability in a bacterial signaling network revealed by single-cell FRET

**DOI:** 10.1101/124008

**Authors:** J.M. Keegstra, K. Kamino, F. Anquez, M.D. Lazova, T. Emonet, T.S. Shimizu

## Abstract

We present *in vivo* single-cell FRET measurements in the *Escherichia coli* chemotaxis system that reveal pervasive signaling variability, both across cells in isogenic populations and within individual cells over time. We quantify cell-to-cell variability of adaptation, ligand response, as well as steady-state output level, and analyze the role of network design in shaping this diversity from gene expression noise. In the absence of changes in gene expression, we find that single cells demonstrate strong temporal fluctuations. We provide evidence that such signaling noise can arise from at least two sources: (i) stochastic activities of adaptation enzymes, and (ii) receptor-kinase dynamics in the absence of adaptation. We demonstrate that under certain conditions, (ii) can generate giant fluctuations that drive signaling activity of the entire cell into a stochastic two-state switching regime. Our findings underscore the importance of molecular noise, arising not only in gene expression but also in protein networks.

## Introduction

Cellular physiology is deeply shaped by molecular fluctuations, resulting in phenotypic variability that can be both detrimental and beneficial (***Rao et al., 2002***). One of the most important and well-studied sources of intracellular fluctuations is stochastic gene expression (***Elowitz et al., 2002; Eldar and Elowitz, 2010; Raj and Van Oudenaarden, 2008***), which can generate substantial cell-to-cell variability in protein levels within isogenic populations under invariant environmental conditions. Such heterogeneity in protein counts are readily measurable by fluorescent-protein reporters (***Elowitz et al., 2002; Ozbudak et al., 2002***), but mechanistically tracing the consequences of such molecular noise to the level of complex cellular phenotypes such as signaling and motility remains a significant challenge, in part due to the multitude of interactions between gene products, but also because each of those interactions can, in principle, become an additional source of noise. In this paper, we study how multiple sources of molecular noise, arising in both gene expression and protein-protein interactions, affect performance of the *E. coli* chemotaxis network, a canonical signaling pathway.

In bacteria, gene-expression noise tends to manifest itself as stable cell-to-cell differences in phenotypes that persist over the cell’s generation time, because typical protein lifetimes are longer than the cell cycle (***Li et al., 2014***). The architecture of signaling networks can have a profound influence on their sensitivity to such noise-induced differences in protein levels, and it has been shown that the design of the *E. coli* chemotaxis network confers robustness of a number of signaling parameters, such as precision of adaptation, against variability in gene expression (***Barkai and Leibler, 1997; Kollmann et al., 2005***). On the other hand, cell-to-cell differences in behavior can also be advantageous for isogenic populations under uncertain and/or time-varying environments, and it has been argued that the manner in which the chemotaxis network filters gene expression noise to shape phenotype distributions could be under selective pressure (***Frankel et al., 2014; Waite et al., 2016***).

In principle, molecular noise arising in processes other than gene expression, such as protein-protein interactions within signaling pathways, can also contribute to cellular variability. However, such noise sources tend to be harder to study experimentally because, in contrast to gene-expression noise, which can be characterized by measuring fluorescent reporter levels (***Elowitz et al., 2002; Raser and O’Shea, 2004***), requirements for *in vivo* measurements of protein-protein interactions tend to be more demanding and no generically applicable strategies exist. The *E. coli* chemotaxis system provides a compelling experimental paradigm for addressing protein-signaling noise, because a powerful technique for *in vivo* measurements of protein signaling, based on Förster resonance energy transfer (FRET), has been successfully developed (***Sourjik and Berg, 2002b; Sourjijk et al., 2007***).

The chemotaxis network controls the motile behavior of *E. coli*, a run-and-tumble random walk that is biased by the signaling network to achieve net migrations toward favorable directions. The molecular mechanisms underlying this pathway have been studied extensively (for recent reviews, see refs. ***Wadhams and Armitage (2004); Tu (2013); Parkinson et al. (2015)***). In brief, transmembrane chemoreceptors bind to ligand molecules, inhibiting the autophosphorylation of a central kinase, CheA. When active, CheA transfers its phosphate to CheY to form CheY-P. Meanwhile, the phosphatase CheZ degrades CheY-P to limit the signal lifetime. CheY-P binds to a flagellar motor, which in turn increases the chance of the motor to turn clockwise, leading to a tumble. An adaptation module consisting of the enzymes CheR and CheB implements negative integral feedback by tuning the sensitivity of the chemoreceptors via reversible covalent modifications that restore the kinase activity (and CheY-P level).

Despite its relative simplicity, this pathway exhibits many interesting network-level functionalities, such as cooperative signal amplification (***Segall et al., 1986; Sourjik and Berg, 2002b; Bray et al., 1998***), sensory adaptation (***Barkai and Leibler, 1997; Alon et al., 1999***) and fold-change detection (***Mesibov et al., 1973; Lazova et al., 2011***), and FRET microscopy has proven extremely powerful in characterizing such signal processing of the chemotaxis pathway, especially in *E. coli* (***Sourjik and Berg, 2002b***, ***2004***; ***Shimizu et al., 2010***; ***Oleksiuk et al., 2011***), but also in Salmonella (***Lazova et al., 2012***) and *B. subtilis* (***Yang et al., 2015***). It has been implemented in various ways (***Sourjik and Berg, 2002b***, ***a***; ***Shimizu et al., 2006***; ***Kentner and Sourjik, 2009***), but most commonly by using CFP and YFP as the FRET donor-acceptor pair, fused to CheY and CheZ, respectively. To date, however, nearly all applications of FRET in the bacterial chemotaxis system have been population-level measurements in which signals from hundreds to thousands of cells are integrated to achieve a high signal-to-noise ratio. A pioneering study applied FRET at the single-cell level to study spatial heterogeneities in CheY-CheZ interactions (***Vaknin and Berg, 2004***), but those measurements were limited to relatively short times due to phototoxicity and bleaching.

By exploring a range of fluorescent proteins as FRET pairs, and improving measurement protocols, we have developed a robust method for single-cell FRET measurements of chemotactic signaling dynamics in single bacteria over extended times. The data reveal extensive cell-to-cell variability, as well as temporal fluctuations that are masked in population-level FRET measurements. In contrast to previous single-cell experiments that relied on measurements of motor output or swimming behavior (***Berg and Brown, 1972; Spudich and Koshland, 1976; Segall et al., 1986; Korobkova et al., 2004; Park et al., 2010; Masson et al., 2012***), FRET alleviates the need to make indirect inferences about intracellular molecular interactions through the highly noisy 2-state switching of the flagellar motor, whose response function can vary over time due to adaptive remodeling (***Yuan et al., 2012***). In a typical experiment, we are able to obtain dozens of (up to ∼100) single-cell FRET time series simultaneously, to efficiently collect statistics of phenotypic and temporal variability.

## Results

### Single-cell FRET reveals pervasive cell-to-cell variability in intracellular signaling

To measure variability in intracellular signaling, we adapted a FRET assay for chemotaxis widely used for population-level measurements with fluorescent fusions to CheY and its phosphatase CheZ (***Sourjik and Berg, 2002b***). On timescales longer than the relaxation of CheY’s phosphorylation/dephosphorylation cycle, the FRET level reflects the phosphorylation rate of CheY by the CheA kinase, thus providing an efficient *in vivo* measurement of the network activity (Fig. 1 - Supplement 1). Instead of the conventional CFP/YFP FRET pair we used the fluorophores YFP and mRFP1 to avoid excitation with blue light, which induces considerably stronger photoxicity and also perturbs the chemotaxis system as a repellent stimulus (***Taylor and Koshland, 1975; Taylor et al., 1979; Wright et al., 2006***). A field of *E. coli* cells expressing this FRET pair were immobilized on a glass surface imaged in two fluorescence channels, and segmented offline to compute a FRET time series for each cell in the field of view (see **Materials and Methods**).

For wild-type cells (Fig. 1a) we found that the ensemble mean of single-cell FRET responses, 〈FRET〉(*t*), agrees well with previous population-level measurements (***Sourjik and Berg, 2002b***). Upon prolonged stimulation with a saturating dose of attractant α-methylaspartate (MeAsp), 〈FRET〉(*t*) rapidly fell to zero before gradually returning to the pre-stimulus level due to adaptation. Upon removal of attractant, 〈FRET〉(*t*) rapidly increased to a maximum before returning to the pre-stimulus baseline. Single-cell FRET time series, FRET*_i_*(*t*), had qualitatively similar profiles, but the kinetics of adaptation and response amplitudes demonstrate differences from cell to cell. For each cell, FRET_*i*_(*t*) is limited by the slow autophosphorylation of CheA and hence is proportional to a[CheA]_T_,_*i*_. (provided [CheY] and [CheZ] are sufficiently high, see Materials and Methods), in which *a* is the activity per kinase and [CheA]T_*i*_. the total amount of kinases part of the receptor-kinase complex. Hence from the FRET_i_.(*t*) the activity per kinase *a* can be readily determined by normalizing each FRET timeseries by its maximum response *a* = FRET_*i*_./FRET_-max_ (Figure 1b). The steady-state activity *a_0*i*_* is then defined as the activity before the addition of attractant. The baseline activity per cell varies from cell-to-cell *a_0_* (quantified by the coefficient of variation, *CV* = 0.22, Fig. 1c). The network activity controls the flagellar motor rotation, and hence this is consistent with the observation that cells in isogenic population exhibit a variable steady-state tumble frequency (***Spudich and Koshland, 1976; Dufour et al., 2016***).

**Figure 1.**
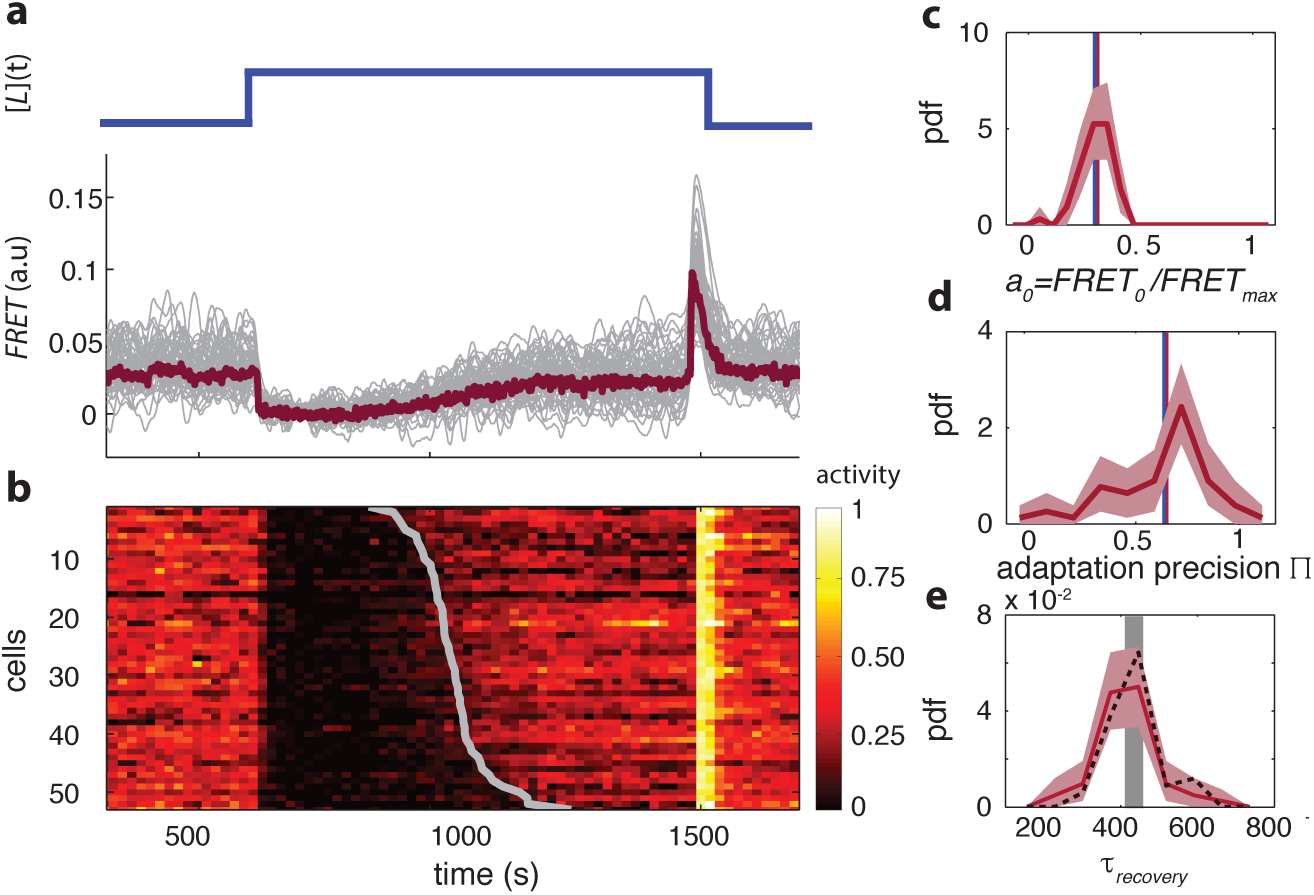
Single-cell FRET over extended times reveals cell-to-cell variability in signaling response. (**a**) Step-response experiment on wildtype cells (CheRB+; VS115). (Top) The ligand time series *[L]*(*t*) indicates the applied temporal protocol for addition and removal of 500 μM MeAsp. (Bottom) FRET response of 54 cells (grey) with the ensemble-averaged time series (dark red) overlaid from a representative single experiment. Single-cell time series were lowpass Filtered with a 14s moving-average Filter. (**b**) Heatmap representation of the normalized FRET response time series, with each row representing a single cell, and successive columns representing the 10s time bins in which the color-indicated activity was computed from the FRET time series. Activity was computed by normalizing FRET to the total response amplitude (Max-Min for each time series). Rows are sorted by the corresponding cell’s recovery time (grey curve), defined as the time at which the activity recovered to 50 % of the activity level after adaptation (see panel e). Single-cell FRET assay schematic and image processing pipeline are shown in Figure 1 - **Supplement 1**. (**c**) Steady-state activity *a*_0_ of the cells shown in panels (a-b). Also shown are the mean steady-state activity (red vertical line) and the steady-state activity of the population averaged time series (blue vertical line). (**d**) Adaptation precision Π obtained from the FRET data. An adaptation precision of 1 denotes perfect adaptation. Also shown are the mean precision (red vertical line) and the precision of the population averaged time series (blue vertical line). The mean and std of the distribution is 0.79 ± 0.32. All colored shaded areas refer to 95 % confidence intervals obtained through bootstrapping. (**e**) Recovery time of cells defined as recovery to 50% of the post-adaptational activity level (red, 54 cells) or 50% of pre-stimulus activity (black dashed, 44 cells with precision >0.5) and simulated effect of experimental noise for a population with identical recovery times (grey). The latter was obtained from a simulated data set in which 55 time series were generated as described in Figure 1 - **Supplement 2**. The width of the bar is defined by the mean plus (minus) the std of the simulated distribution. The mean and std of the distributions for the experimental and simulated data sets are respectively 416 ± 83 and 420 ± 35. **Figure 1 - Supplement 1** Single-cell FRET assay schematic and workflow. **Figure 1 - Supplement 2** Influence of experimental noise on estimating recovery times.

The adaptation precision is defined as its post-adaptational activity level (Π = *a_adapted,i_/a_0,*i*_*), hence a precision of 1 refers to perfect adaptation. The adaptation kinetics are quantified by the recovery time *τ_recovery_*, the time required for each cell to recover to 50% of its post-adaptational activity level (*a*_adapted,*i*_). When observing the distributions of these parameters we noted that the cell-to-cell variability is high in the precision Π (Fig. 1d, *CV*=0.40) but the average precision (0.79) agrees well with population measurements (***Neumann et al., 2014***). The variation is also substantial in *τ_recovery_*(Fig. 1d, *CV*=0.20). This falls within the range of ∼20-50% from previous reports, in which single-cell recovery times were estimated from motor-rotation or swimming-behavior measurements (***Berg and Tedesco, 1975; Spudich and Koshland, 1976; Min et al., 2012).*** The time required to recover from a saturating amount of attractant is determined not only by the stimulus size, but also the methylation rate of receptor modification sites catalyzed by CheR and the number of such sites that need to be methylated. Variability in the recovery time is thus likely to reflect cell-to-cell variability in the ratio between the expression level of CheR and that of the chemoreceptor species responding to ligand (Tar for the experiment in Fig. 1a).

Thus, single-cell FRET allows efficient measurement of single-cell signaling dynamics that, on average, agree well with previous population-level FRET experiments and single-cell flagellar-based experiments, thereby revealing variability in a wide variety of signaling parameters.

### Diversity in the ligand response is modulated during population growth

The chemoreceptor clusters in *E. coli* are the central processing units and are responsible for signal integration and amplification. The sensory output of the cluster, the activity of the kinase CheA, is activated by a mixture of chemoreceptors. Cooperative interactions within the kinase-receptor complex leads to amplifications of small input stimuli and weighting different input signals. It has been shown that the composition of the receptor-kinase complexes can affect both the amplification as well as the weighting of different input signals (***Ames et al., 2002; Sourjik and Berg, 2004; Kalinin et al., 2010***), but how the amplification and integration varies across a population has not been characterized. To bridge the gap between collective behavior and its underlying single-cell motility it is essential to determine the variability of these important signaling parameters, as well as the origin of the variability. Also, current estimates of the apparent gain in the response (defined as the fractional change in output divided by fractional change in input) are based on population-averaged measurements which may may not reflect single-cell cooperativity levels. In population averaged measurements, the largest gain is observed in adaptation-deficient (CheRB-) cells (***Sourjik and Berg, 2004***), in which the receptor population is homogeneous with respect to their adaptational modification state and hence in these cells variability in ligand sensing can be studied separately from variability induced by the adaptation enzymes.

We probed the ligand sensitivity of CheRB-cells (TSS58) at the single-cell level by FRET dose-response measurements in which step stimuli of successively larger amplitudes were applied over time (Fig. 2). Considerable variability in the response to the attractant L-serine were observed across the population of immobilized cells simultaneously experiencing the same stimulus, with response magnitudes often ranging from virtually zero to full response (Fig. 2a). The resulting dose-response data were analyzed by fitting each individual cell’s FRET response by a Hill curve of the form [1 + ([*L*]/*K*)^H^]^−1^, where the parameters (1/*K*) and *H* are defined as the sensitivity and steepness, respectively, of the response. The family of dose response curves constructed from this ensemble of fit parameters reveals considerable variability from cell to cell in the shape of the response curve (Fig. 2b).

**Figure 2.**
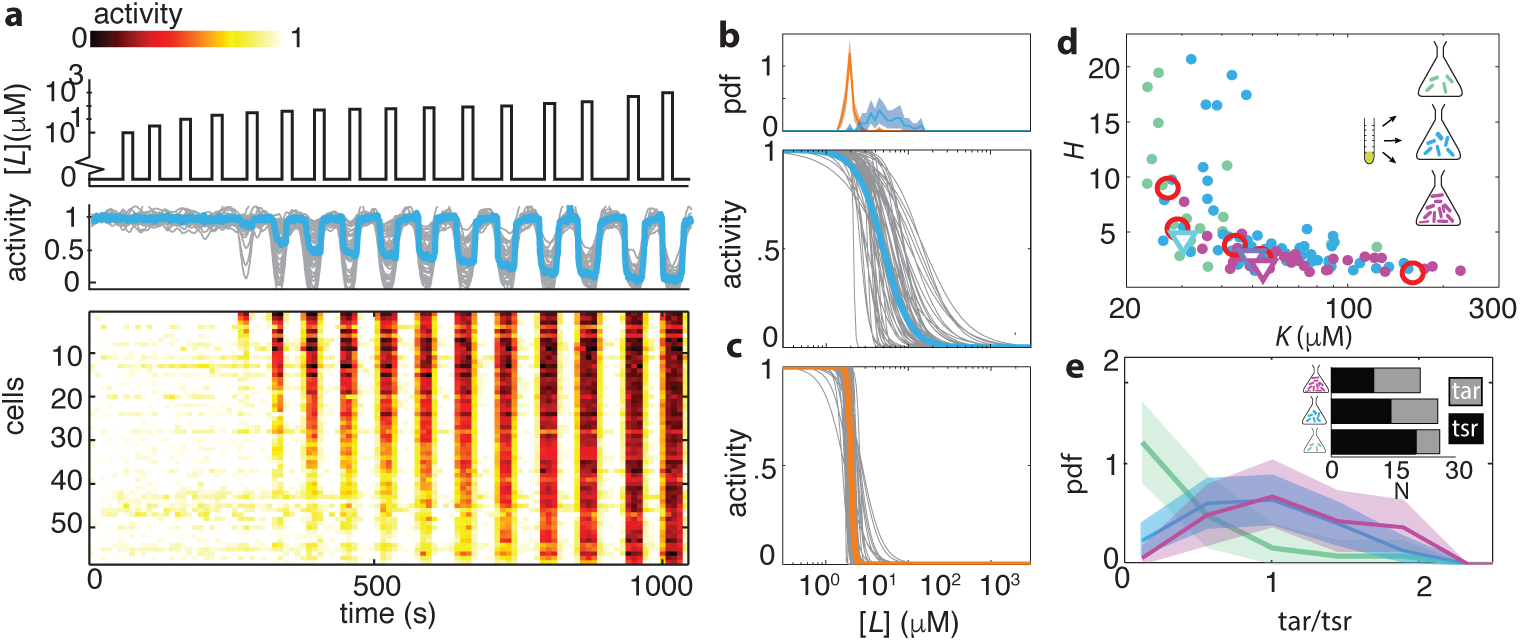
Ligand dose-response parameters vary strongly across cells in an isogenic population, even in the absence of adaptation, and depend on receptor-complex composition. (**a**) Single-cell dose-response experiment on adaptation deficient (CheRB-; TSS58) cells with a wildtype complement of receptors. (Top) Temporal protocol of stimulation *[L]*(*t*) by the attractant L-serine. (Middle) The ensemble-averaged FRET response of the population (blue) and single cells (gray) in signaling activity of 59 cells from a single experiment, normalized to the full-scale FRET response amplitude. (Bottom) Heatmap representation of the single-cell FRET timeseries, with the rows sorted by the sensitivity *K* of the corresponding cell obtained from Hill-curve fits. (**b**) Family of dose response curves (gray) obtained from the Hill-curve fits to single-cell dose-response data. CheRB-cells with a wildtype complement of receptors (TSS58). The blue curve was obtained from fitting a hill function to the population-averaged time series shown in panel (**a**), yielding fit values*K*=50 μM and *H*=2.7. The fitted single-cell *K* values are shown in the histogram on top (blue). (**c**) CheRB-cells expressing only the serine receptor Tsr (UU2567). The orange curve was obtained from fits to the population average, yielding K=20 μM and H=22. The fitted single-cell *K* values are shown in the histogram in panel (b) (orange). (**d**) Cells from a single overnight culture were innoculated into three flasks harvested at different times during batch-culture growth growth to sample the state of the population at three points along the growth curve: at OD_600_ = 0.31 (green), 0.45 (blue) and 0.59 (purple). The fits to the population-averaged time series are shown in Figure 2 - **Supplement 1**. Shown are Hill-curve sensitivity (1/K) and cooperativity *H* obtained from fits to the single-cell time series, at different OD’s (filled dots) together with the fit values from the population-average (triangles). Also shown are population-FRET results in which Tar and Tsr levels were controlled artificially (red *circles*,(*Sourjik and Berg, 2004*)). Shown are 25 out of 28 cells harvested at OD=0.31, 59 out of 64 cells at OD=0.45, 34 out of 40 cells at OD=0.59. The excluded cells had fits with a mean squared error higher then 0.05. The influence of experimental noise on the fit parameters is shown in Figure 2 - **Supplement 2**. **e**) Histograms of Tar/Tsr ratio obtained from fit of multi-species MWC model from reference **Mello and Tu (2005)** to single-cell FRET time series. The mean Tar/Tsr ratios (low to high OD) are 0.4, 0.9, and 1.2 with coefficients of variance of respectively 1.1, 0.5, and 0.4. Inset: average cluster size of Tar (grey) and Tsr (black) at different harvesting OD’s obtained from the fit results in panel d. Figure 2 - Supplement 1 Dose response curves from population averaged time series at different harvesting OD’s. Figure 2 - Supplement 2 Influence of experimental noise on fit parameters *K* and *H*.

What could be the cause of the diversity in ligand response in the absence of adaptation-induced heterogeneity? We reasoned that expression-level variability of the five chemoreceptor species of *E. coli*, which are known to form mixed clusters with cooperative interactions (***Ames et al., 2002; Sourjik and Berg, 2004***), could endow isogenic populations with sensory diversity. In line with this idea, CheRB-cells expressing only a single chemoreceptor species (Tsr) demonstrated not only higher cooperativity, but also attenuated variability in the dose-response profile from cell to cell (Figure 2b-c), showing that the composition of the receptor population is important not only to tune the average ligand response of a population, but also in generating a wide range of sensory phenotypes within an isogenic population.

It has been shown that expression level of chemoreceptors changes during growth of *E. coli* batch cultures: concomitant with the slowing of growth upon the transition from the exponential phase towards early stationary phase, the relative expression level ratio Tar/Tsr, the two most abundant chemoreceptors, increases from majority Tsr (Tar/Tsr< 1) to majority Tar (Tar/Tsr> 1) (***Salman and Libchaber, 2007***; ***Kalinin et al., 2010***). To probe the consequence of such changes for ligand-sensing diversity, we measured single-cell dose response curves in populations harvested at different cell densities during batch growth (Figure 2d). The resulting population-averaged responses show a dependence of dose-response parameters on the optical density (O.D.) of the culture, shifting from highly sensitive (low *K*) and highly cooperative (high *H*) at low cell densities (OD≈ 0.3) to less sensitive (high *K*) and less cooperative (low *H*) at increased cell densities (OD≈ 0.45, and OD ≈ 0.6) (Fig. 2d, open triangles, and Fig. 2 - Supplement 1). This trend is also visible at the level of single cells, but we found the responses to be highly variable under each condition (Fig. 2d, filled points). Remarkably, both *K* and *H* varied by over an order of magnitude, far exceeding the uncertainty in parameter estimates due to experimental noise (Fig. 2 - Supplement 2).

To further test the idea that ligand-response diversity is governed by differences in receptor expression levels, we considered the pattern of covariation between the fitted sensitivity *K* and cooperativity *H* in single cells (Figure 2b, blue). In contrast to cells expressing Tsr as the only chemoreceptor, in which the variability in *K* is only 20 % (Figure 2b, orange), single cells expressing a wildtype complement of chemoreceptors demonstrated strong variation in *K*. This variation was negatively correlated with the cooperativity *H* (Figure 2d). Noting that this overall pattern of covariation agrees well with dose response parameters obtained from population-level FRET experiments in which the Tar/Tsr ratio was experimentally manipulated via plasmid-based expression control (Figure 2d, open triangles; data from ***Sourjik and Berg (2004)***), we proceeded to quantitatively estimate the diversity in the Tar/Tsr ratio via fits of a multi-species MWC model (***Mello and Tu, 2005***; ***Keymer et al., 2006***) to single-cell FRET data (see Materials and Methods). The resulting distribution of single-cell Tar/Tsr estimates (Figure 2e) was dominated by Tsr in cells harvested early (OD≈ 0.3) but the relative contribution of Tar increased in cells harvested at later stages of growth (OD≈ 0.45 and OD≈ 0.6). Interestingly, in addition to this increase in the mean of the Tar/Tsr distribution during batch growth, which confirms previous reports that found increased Tar/Tsr ratios at the population level (***Salman and Libchaber, 2007***; ***Kalinin et al., 2010***), we find that the breadth of the distribution also increases at later stages of growth. Thus, modulation of receptor expression during growth provides a means of tuning not only response sensitivity and cooperativity, but also single-cell diversity in the response of cell populations experiencing identical changes in their common environment.

The large variability in the Tar/Tsr ratio (*CV* ≈ 0.5 at O.D.=0.45) is somewhat surprising given that the mean expression level of both receptors are known to be high and of order 10^3^-10^4^ copies per cell (***Li and Hazelbauer, 2004***). At such high expression, intrinsic noise in expression levels (i.e. due to the production and degradation process of proteins, expected to scale as the square root of the mean) could be as low as a few percent of the mean, and gene-expression fluctuations are expected to be dominated by extrinsic noise components (i.e. those affecting regulation of gene expression, which do not scale with the mean). In *E. coli*, a global survey of gene expression noise established an empirical lower bound to extrinsic noise at *CV* ≈ 0.3 (***Taniguchi et al., 2010***), and measurements within the chemotaxis system also indicate that between a subset of chemotaxis genes, the extrinsic component of covariation does approach this limit (***Kollmann et al., 2005***). Interestingly, however, a recent study (**Yoney and Salman, 2015)** found using single-cell flow-cytometry a high degree of variability in the ratio of Tar/Tsr promotor activities (*CV* ≈ 0.45 at O.D.=0.51) comparable to the range of ratios extracted from our analysis of dose response data. Given that cell-to-cell variability in the Tar/Tsr ratio is much greater than achievable lower bounds of gene-expression noise in bacteria, it would be interesting to investigate the mechanistic sources of this variability, such as operon organization, RBS strength, and promotor stochasticity (***Frankel et al., 2014***).

Variability in receptor expression could also explain the distribution of adaptation precision we observed in wildtype cells (Figure 1d). In a previous population-level study, it has been shown that adaptation precision depends strongly on the expression-level ratio between the multiple chemoreceptor species, with the highest adaptation precision being achieved when the ligand-binding receptor is a minority within the total receptor population (***Neumann et al., 2014***). Thus, the substantial heterogeneity in adaptation precision we observed (*CV* = 0.40) upon a saturating MeAsp stimulus is consistent with strong variability in the Tar/Tsr ratio.

### CheB phosphorylation feedback attenuates cell-to-cell variability

While bacteria can exploit molecular noise for beneficial diversification, variability can also limit reliable information transfer and degrade sensory performance. In the framework of *E. colis* run- and-tumble navigation strategy, chemotactic response to gradients requires that cells maintain a finite tumble bias, the fraction of time a bacterium spends tumbling, and avoids extreme values zero and one. The latter cases would correspond to unresponsive phenotypes that fail to switch between run and tumble states in response to the environmental inputs. One important mechanism that ensures responsiveness to stimuli over a broad range of input levels is sensory adaptation mediated by the methyltransferase/methylesterase pair CheR/CheB. These receptor-modifying enzymes provide negative feedback through the dependence of their catalytic activity on the receptor’s signaling state: the rate of methylation (demethylation) by CheR (CheB) is a decreasing (increasing) function of receptor-kinase activity (***Borczuk et al., 1986; Amin and Hazelbauer, 2010***).This dependence of enzyme activity on the substrate conformation provides negative integral feedback that ensures precise adaptation (***Barkai and Leibler, 1997***) toward the pre-stimulus steady-state activity *a*_0_.

Interestingly, one of the two adaptation enzymes, CheB, can be phosphorylated by CheA, the kinase whose activity CheB controls through its catalytic (demethylation) activity on receptors. Effectively, this adds an additional negative feedback loop to the network, but the role of this phosphorylation-dependent feedback has remained elusive since it has been shown to be dispensable for precise adaptation (***Alon et al., 1999***). Through theoretical analysis, it has been conjectured that this secondary feedback loop might play a role in attenuating effects of gene-expression noise (***Kollmann et al., 2005***), but experimental verification has been lacking. We therefore sought to investigate the influence of perturbations to this network topology on the variability of chemotactic signaling activity.

CheB consists of two domains connected by a flexible linker (Figure 3a). A regulatory domain, with structural similarity to CheY, can be phosphorylated at residue Asp^56^ (***Djordevic et al., 1998; Stewart et al., 1990***). A catalytic domain mediates binding to specific residues on chemoreceptor cytoplasmic domains and removes a methyl group added by the counterbalancing activity of CheR. Phosphorylation induces a conformational change and activates CheB (CheB*) (***Djordevic et al., 1998; Lupas and Stock, 1989***). Several mutants of CheB lack phosphorylation feedback while retaining catalytic activity. Here, we focus on two specific mutants: CheB^D56E^, which bears a point mutation at the phosphorylation site, and CheB_c_, which expresses only the catalytic domain of CheB (***Stewart et al., 1990; Alon et al., 1999***). Cells expressing these mutants have an altered network topology (Figure 3b) which lacks CheB phosphorylation feedback.

**Figure 3.**
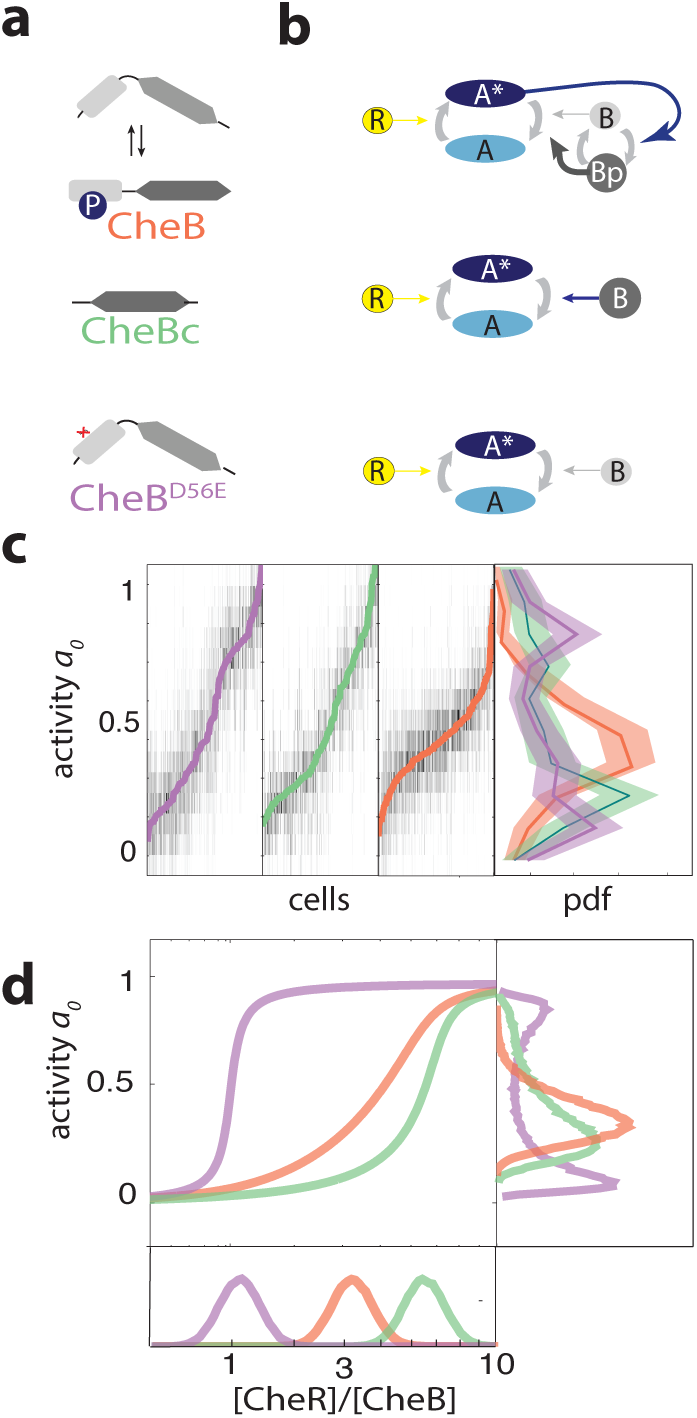
CheB phosphorylation feedback attenuates variability in steady-state kinase activity. (**a**) Schematic depiction of CheB activation by phosphorylation. (Top) CheB consists of two domains connected by a flexible linker. The aspartate at residue 56 can be phosphorylated. (Middle) CheBc lacks the receiver domain with the phosphorylation site. (Bottom) CheB-D56E carries a point mutation at the phosphorylation site. (**b**) Effective network topology of cells expressing WT CheB (top), CheBc (middle) and CheB-D56E (bottom). All network topologies can have perfect adaptation. Using population FRET, we found that phosphorylation feedback is not required for fast removal adaptation kinetics (Figure 3 - **Supplement 3**). (**c**) Heatmap representation of histograms of activity *a*(*t*) around steady-state of single-cells, from experiments as shown in Figure 3 - **Supplement 1**. Each colum represents a single cell, for each CheB mutant in a *cheB* background (VS124, colors as in panel (a)). The single cells are ordered by the steady-state activity *a*_0_, which is superimposed. (right) Histograms with *a_0_* for each CheB mutant. All histograms contain the results for cells with a signal-to-noise ratio higher than 1 from at least 3 independent FRET experiments, which corresponds to 322 out of 373 cells (WT), 225 out of 279 cells (CheBc) and 226 out of 359 cells (D56E). Shaded regions represent bootstrapped 95% confidence intervals. We verified that the bistability was caused by clipping of extreme values due to potential FRET pair saturation, by working in a regime where the response amplitude cannot be saturated by the FRET pair (Figure 3 - **Supplement 2**). Furthermore, we established that the defect in phosphorylation leads to impared chemotaxis in soft agar plates (Figure 3 - **Supplement 4**). (**d**) A simple kinetic model of the chemotaxis network illustrates the crucial role of CheB phosphorylation feedback in circumventing detrimental bimodality in *a*_0_. Due to saturated enzyme kinetics in the adaptation system, the transfer function between [CheR]/[CheB] expression ratio and steady-state network output *a_0_* can be highly nonlinear (main panel). The shape of this transfer function determines the distribution of *a_0_* (right panel) by transforming the distribution of [CheR]/[CheB] expression ratios (bottom panel). Shown are a CheB version with lower activity as WT (purple) and higher than WT (green), both without feedback, and one CheB species with phosphorylation feedback (orange). **Figure 3 - Supplement 3** Phosphorylation feedback is not a necessary condition for fast removal adaptation dynamics. **Figure 3 - Supplement 1** Example FRET time series and CheB localization. **Figure 3 - Supplement 2** Relation between maximum FRET response and FRET fluorophore expression levels. **Figure 3 Supplement 4** Phosphorylation defective mutants show impaired chemotaxis on soft agar.

To study the influence of network topology on cell-to-cell variability, we expressed different forms of CheB (CheB^WT^, CheB^D56E^, CheB_c_) from an inducible promoter in a *ΔcheB* strain and measured the response to a saturating amount of attractant (500 μM MeAsp). The expression levels of each mutant are tuned such that they approximate the wild-type steady state activity level. The response variability of CheB^WT^ was, as expected, very similar to cells in which CheB is expressed from its native chromosomal position (compare Fig. 3 - Supplement 1a and Fig. 1a). By contrast, cells expressing either of the two CheB mutants defective in phosphorylation demonstrated increased cell-to-cell variability in the steady-state activity compared to cells expressing CheB^WT^. The increased variability of the CheB phosphorylation-deficient mutants (CheB^D56E^ and CheB_c_)was manifested not only in a higher coefficient of variation in *a_0_* (1.07 and 1.10, respectively, and WT 0.7), but also a qualitatively different shape of the distribution of *a_0_* across the population (Figure 3c). Whereas the distribution demonstrated a single peak in CheB^WT^ cells with phosphorylation feedback, the distribution for the phosphorylation-feedback mutants demonstrated a bimodal shape with peaks close to the extreme values *a_0_* = {0,1}.

We tested whether these strong differences in cell-to-cell variability might be the result of gene expression noise, by comparing expression-level distributions of the CheB mutants. We constructed fluorescent fusions of each *cheB* allele to the yellow fluorescent protein mVenus and quantified the distribution of single-cell fluorescence levels under the same induction conditions as in the FRET experiments (figure 3 - Supplement 1). The ratio between the measured expression-levels (CheBc:WT:D56E≈0.7:1:2.5) was compatible with expectations from the hierarchy of reported *in vitro* catalytic rates of CheB 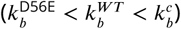(***Anand and Stock, 2002; Simms et al., 1985; Stewart, 1993***), and expression-level variability was very similar between the three strains (*CV*’s of 0.87,0.90 and 0.82). These findings suggest that the differences in cell-to-cell variability observed in FRET are not due to differences between the expression-level distributions of the three *cheB* alleles, but rather to the differences they impose on the signaling network topology.

What feature of the signaling network could generate such broad (and even bimodal) distributions of *a*_0_? It has been conjectured (***Barkai and Leibler, 1997; Emonet and Cluzel, 2008***) and demonstrated (***Shimizu et al., 2010***) that *in vivo* the enzymes CheR and CheB operate at or near saturation. An important consequence of enzyme saturation in such reversible modification cycles is that the steady-state activity of the substrate becomes highly sensitive to the expression level ratio of the two enzymes, a phenomenon known as zero-order ultrasensitivity (***Goldbeter and Koshland (1981);*** see **Materials and Methods**). Within the chemotaxis system, saturation of both CheR and CheB can thus render the receptor modification level, and in turn, the CheA activity *a_0_*, ultrasensitive to the [CheR]/[CheB] concentration ratio. If we view this ultrasensitive mapping as a transfer function *f* between the ratio [CheR]/[CheB] and the steady-state activity *a*_0_, 
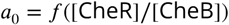

then its characteristic steep sigmoidal profile can impose bimodality in the methylation level, and hence also in the activity of steady-state CheA activity, *a*_0_, even at quite modest input variances for distributions of the ratio *P_RB_*([CheR]/[CheB]). This is because the manner in which the transfer function *f* filters the [CheR]/[CheB] distribution, 
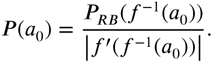
spreads the narrow range in the [CheR]/[CheB] distribution *P_RB_*([CheR]/[CheB]) over which *f* ([CheR]/[CheB]) is steep across a broad range in *a_0_.* Thus, even if expression-level noise for both CheR and CheB are modest, an ultrasensitive transfer function *f* can effectively amplify the variation in [CheR]/[CheB], and if the distribution of the latter ratio,*P_RB_*([CheR]/[CheB]) extends below and above the narrow region over which *f* is steep, the decreased slope of *f* (i.e. lower *f*’([CheR]/[CheB])) in those flanking regions will tend to increase the weight on both sides of the broad *P*(a_0_) distribution to produce a bimodal profile. On the other hand, if the network topology effectively reduces the steepness of *f*, the resulting *P*(*a*_0_) will have a reduced variance for the same input *P_RB_*([CheR]/[CheB]) (Figure 3d).

Could the known biochemical differences between the three forms of CheB (CheB^WT^, CheB^D56E^, CheB_c_) explain the contrasting patterns of *a*_0_ variability observed in our single-cell FRET experiments? In the absence of any feedback, the steepness of *f (CheR/CheB*) is solely determined by the low Michaelis-Menten constants *K_B R_*, which corresponds to saturated kinetics of the enzymatic activity of CheRB and hence ultransensitivity of the steady-state substrate activity. The expression ratio of CheR/CheB which determines the crossover point (*a*_0_=0.5) is set by the ratio of catalytic rates of CheR and CheB (*k*_rb_). Hence the phosphorylation deficient mutants CheB^D56E^ and CheB_c_ both have steep curves but are shifted along the R/B axis due to very different catalytic rates. However,in the case of phosphorylation feedback, CheB^WT^, the same enzyme can be in two states, each with equal *K_rb_* but one low and one high *k_r_.* Whether CheB is in the one state or the other is determined by the activity-dependent phosphorylation feedback. As a result, the curve of CheB^WT^ is activity dependent (f (a,*CheR/CheB*)) and changes with activity by shifting between the two curves corresponding to the extremes of all phosphorylated or all unphosphorylated. Effectively, this makes the resulting curve *f* less steep. The mean of the distributions *P_RB_* are tuned such to get the same mean activity level (〈a_0_〉), but the same variance in *P_RB_* leads to very wide *P*(*a*_0_) distributions in absence of phosphorylation, while phosphorylation feedback ensures a much smaller, single-peaked distrubtion.

It has also been conjectured that the CheB phosphorylation feedback is responsible for the highly nonlinear kinetics of recovery from repellent (or attractant removal) responses ***Shimizu et al. (2010***). Indeed, in cells expressing CheB_c_, the kinetics of recovery from the response to removal of 500 μM MeAsp after adaptation appeared qualitatively different from that in cells expressing wildtype CheB, lacking the characteristic rapid recovery and instead appearing more symmetric with the CheR-mediated recovery upon addition of a saturating dose of attractant (Fig. 3 - Supplement 3). By contrast, CheB^D56E^ was found to still possesses a fast component, despite being defective in phosphorylation, albeit also with somewhat slower kinetics than wt. In summary, the clearest difference between wildtype and phosphorylation-defective CheB mutants is found in the variability of the steady-state signal output (i.e. kinase activity).

The bimodal distribution in kinase activity we observed in the phosphorylation-deficient CheB mutants implies that a large fraction of cells have a CheY-P concentration far below or far above the motor’s response threhold and hence will impair chemotactic responses to environmental gradients. Consistent with this idea, in motility-plate experiments (Supplementary Figure 3 - **Supplement 4**) we found that chemotactic migration on soft-agar plates was severely compromised for both CheB^D56E^ and CheB_c_ compared to CheB_WT_, indicating that the phosphorylation feedback is important for efficient collective motility.

### Protein-signaling noise generates large temporal fluctuations in network output

The slow kinetics of the adaptation enzymes CheR and CheB have been hypothesized to play a role not only in determining the steady-state kinase activity *a_0_*, but also in generating temporal fluctuations of the intracellular signal (***Korobkova et al., 2004; Emonet and Cluzel, 2008; Park et al., 2010; Celani and Vergassola, 2012***). We found substantial differences between wildtype (CheRB+) and adaptation-deficient (CheRB-) cells in the variability of their FRET signals across time (Fig. 4). The effect is clearly visible upon comparing long (∼1h) FRET time series obtained from cells of these two genotypes (Figure 4a). The FRET signal in wildtype cells demonstrated transient excursions from the mean level that were far greater in amplitude than those in CheRB-cells. This amplitude *η* ≡ *σ_a_/*〈*a*〉 was quantified by computing the variance of each single-cell time series, low-pass filtered with a moving average filter of 10s, and shows that the fluctuation amplitudes are much larger in wildtype cells compared to adaptation-deficient cells ((η) =0.09 and 0.44, Fig. 4b). Importantly, these experiments were carried out under conditions in which no protein synthesis can occur due to auxotrophic limiation (see Materials and Methods), thus ruling out gene-expression processes as the source of these fluctuations.

**Figure 4.**
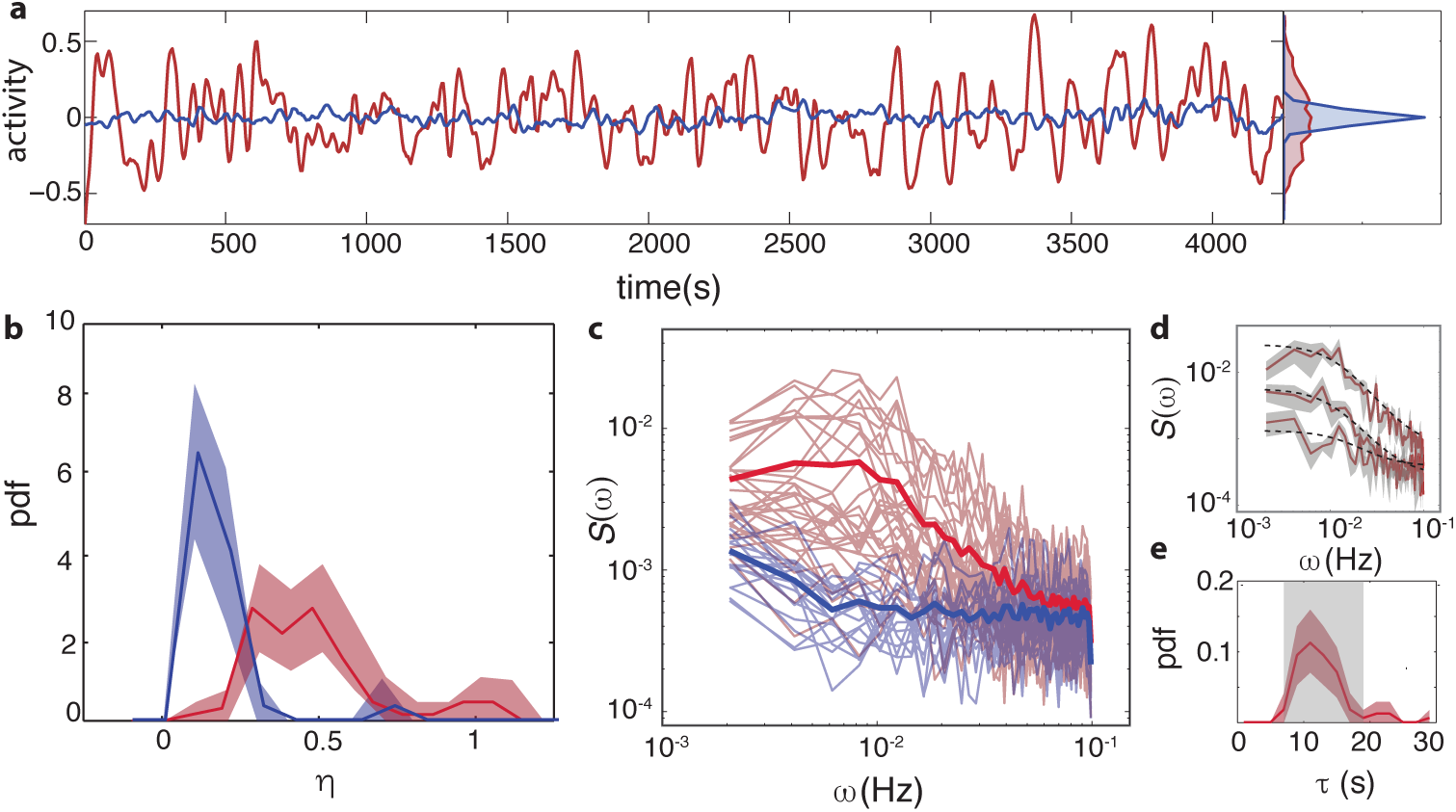
Temporal signal fluctuations in the absence of ligand stimulation are generated by stochastic activity of the adaptation enzymes CheR/CheB. (**a**) Representative single-cell FRET time series of steady-state fluctuations in the presence (CheRB+, VS115, red) and absence (RB-,TSS58,blue) of adaptation kinetics in normalized activity units. (**b**) Histogram of fluctuation amplitude *η* for both CheRB+ (89 cells, red, from 3 independent experiments) and CheRB- (33 cells, blue, from two independent experiments), extracted from calculating the standard deviation of a low-pass Filtered FRET time series over a 10s window divided by the mean FRET level of a single cell. The shaded areas represents the 95% conAdence interval obtained from bootstrapping. (**c**) Power spectral density (PSD) computed from single-cell FRET time series of 31 CheRB+ cells (red, from single experiment) and 17 CheRB-cells (blue, from single experiment), each from a single experiment. Thin curves in the lighter shade of each color represent Single-cell spectra, and the thick curves in the darker shade are the average of all single-cell spectra for each genotype. Note that if the PSD is calculated from the population average the effect of fluctuations are lost Figure 4 - **Supplement 1**. (**d**) Representative single-cell PSDs and fits by an Ornstein-Uhlenbeck (O-U) process. Shown are O-U Fits (Lorentzian with constant noise floor; dashed curves) to three single-cell PSDs (solid curves). The shaded area represents the standard error of the mean for PSDs computed from nine non-overlapping segments of each single-cell time series. Fits to all cells from the same experiment are shown in (Figure 4 - **Supplement 2**). From the OU Fit parameters the noise amplitude can be calculated (Figure 4 - **Supplement 3**). (**e**) Histogram of fluctuation timescales *τ* extracted from single-cell PSD Fits (red, 75 out of 89 cells). Cells without a clear noise plateau were excluded from the analysis (Figure 4 - **Supplement 3**). The shaded areas represents the 95% conAdence interval obtained from bootstrapping. The grey shaded area refers to the variability (mean±std) that can be explained by experimental noise and a finite time window, obtained through simulated O-U time series (see **Materials and Methods**). **Figure 4 - Supplement 1** PSD estimates from population-averaged time series. **Figure 4 - Supplement 2** Fits of OU process to PSD estimates from single-cell FRET time series from a representative single experiment. **Figure 4 - Supplement 3** Comparison between noise amplitudes obtained from time series and power spectra.

Power spectral density (PSD) estimates computed from such time series confirm a nearly flat noise spectrum for CheRB-cells, whereas CheRB+ cells demonstrated elevated noise at low frequencies (Fig. 4c). The amplitude of these low-frequency noise components do clearly vary from cell to cell, as can be gleaned in the diversity of single-cell power spectra. To quantify this protein-level noise due to CheR/CheB activity, we describe the fluctuating signal as an Ornstein-Uhlenbeck (O-U) process, with relaxation timescale ***τ*** and diffusion constant c, which can be interpreted as a linear-noise approximation (***Van Kampen, 1981***) to the full stochastic chemical kinetics of the network controlling the mean kinase activity ***a (Emonet and Cluzel, 2008):***

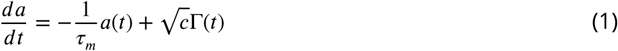
where Γ(*t*) is a Gaussian white noise process. The parameters *τ_Μ_* and *c* for each cell are readily extracted via the power-spectrum solution of the O-U process: 
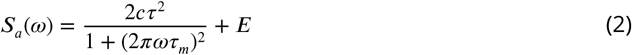
where we have added to the standard Lorentzian solution (***Gillespie, 1996***) a white-noise term *E* that may vary from cell to cell to account for experimental shot noise in the photon-limited FRET signal. Single-cell PSD data were well fit by Eq. 2 (Figure 4d), and the average of extracted single-cell fluctuation timescales (τ_m_〉 = 12.6s) (Figure 4e) are in good agreement with previously reported correlation times of flagellar motor switching (***Park et al., 2010***; ***Korobkova et al., 2004***), as well as the kinetics of CheRB-mediated changes in receptor modification from *in vivo* measurements using radioactively labeled methyl groups (***Lupas and Stock, 1989; Terwilliger et al., 1986***). The variance of the fluctuations obtained from the fits of the PSD, 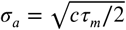 yielded very similar noise amplitudes *η_OU_* ≡ *σ_a,OU_*/〈*a*〉 as calculated from the time series (〈*η_OU_*〉 ≡ 0.42, Fig. 4 - Supplement 3). We note that these noise levels are larger than expected - in a considerable fraction of cells, the standard deviation of fluctuations is comparable to the mean level of activity, and the steady-state fluctuations span the full range of kinase activity (see e.g. that represented by the red curve in Fig. reffig:results4a). Previous studies had predicted a value of ∼ 10 – 20%, based either on reported fluctuation amplitudes of motor switching (***Korobkova et al., 2004; Tu and Grinstein, 2005***)or biochemical parameters of the intracellular signaling network (***Emonet and Cluzel, 2008; Shimizu et al., 2010***) and is also highly variable (σ_*η*_ = 0.24) from cell to cell.

In summary, we confirmed the presence of strong temporal fluctuations in single-cell chemo-taxis signaling attributable to the stochastic kinetics of the adaptation enzymes CheR/CheB, and further found that the amplitude of these fluctuations vary considerably across cells in an isogenic population.

### Receptor-kinase fluctuations in the absence of adaptation reveal two-level switching

The fluctuation amplitude *η* in CheRB+ cells (Fig. 4b) is much greater than previous estimates from pathway-based models that considered zero-order ultrasensitivity in the enzymatic activities of CheR and CheB (***Emonet and Cluzel, 2008***) and receptor cooperativity (***Shimizu et al., 2010***) as possible mechanisms that amplify noise originating in the stochastic kinetics of receptor methylation/demethylation. A plausible explanation for the latter discrepancy is that receptor cooperativity, which can amplify not only ligand signals but also fluctuations in receptor methylation levels (***Duke and Bray, 1999; Shimizu et al., 2003; Mello et al., 2004***), is actually much stronger at the single-cell level than was previously estimated from population-level FRET measurements. The dose-response data for CheRB-cells presented in this study (Fig. 2) clearly demonstrate that single-cell dose-response curves tend to be steeper than those obtained from fits to the population average. Yet another (though not exclusive) possibility is that there exist significant sources of signaling noise that are independent of enzymatic receptor modification. Although we found that the noise amplitude *η* was much lower than wildtype in unstimulated CheRB-cells (Fig. 4), if the cooperative switch-like signaling response of these cells to chemoattractant stimulation (Fig. 2) apply also to other perturbations, it is possible that the strong activity bias in the absence of chemoeffectors (*a*_0_ ≈ 1) masks noise contributions that would be observable if receptors were tuned to the more responsive regime of intermediate activity. In wildtype cells, such a tuning is achieved by the balance between CheR and CheB activities, which sets the steady-state receptor-kinase activity to an intermediate value (*a*_0_ ≈ 0.2-0.5; see Fig. 1).

We reasoned that in CheRB-cells, tuning the activity to an intermediate level by adding and sustaining a sub-saturating dose of attractant could reveal additional noise sources, if present and of significant strength. To maximize detection sensitivity, we focused on cells expressing Tsr as the sole chemoreceptor in a CheRB-background, which demonstrate the highest cooperativity in ligand responses and hence could be expected to strongly amplify also noise sources relevant to signaling (Fig. 2b). To test the effect of bringing the system to the responsive regime, we applied first a large saturating dose of Tsr’s cognate ligand L-serine, followed by a prolonged stimulus of a magnitude close to the dose-response parameter *K*, eliciting a half-maximal population-level response (Fig. 5a). During the second stimulus, which was sustained for several minutes, the population-level response remained approximately constant. Strikingly, however, the time series of single-cell responses demonstrated strong deviations from the population average (Fig. 5b). Whereas all cells responded identically to the saturating dose of attractant, the behavior during the sub-saturating step was highly diverse. Some cells (11/141) showed no apparent response in kinase activity, whereas in others (32/141) complete inhibition was observed (Fig. 5b, yellow curves). The majority cells (98/141), however, had an intermediate level of activity when averaged over time, but demonstrated strong temporal fluctuations, often with magnitudes exceeding those observed in wildtype cells.

**Figure 5.**
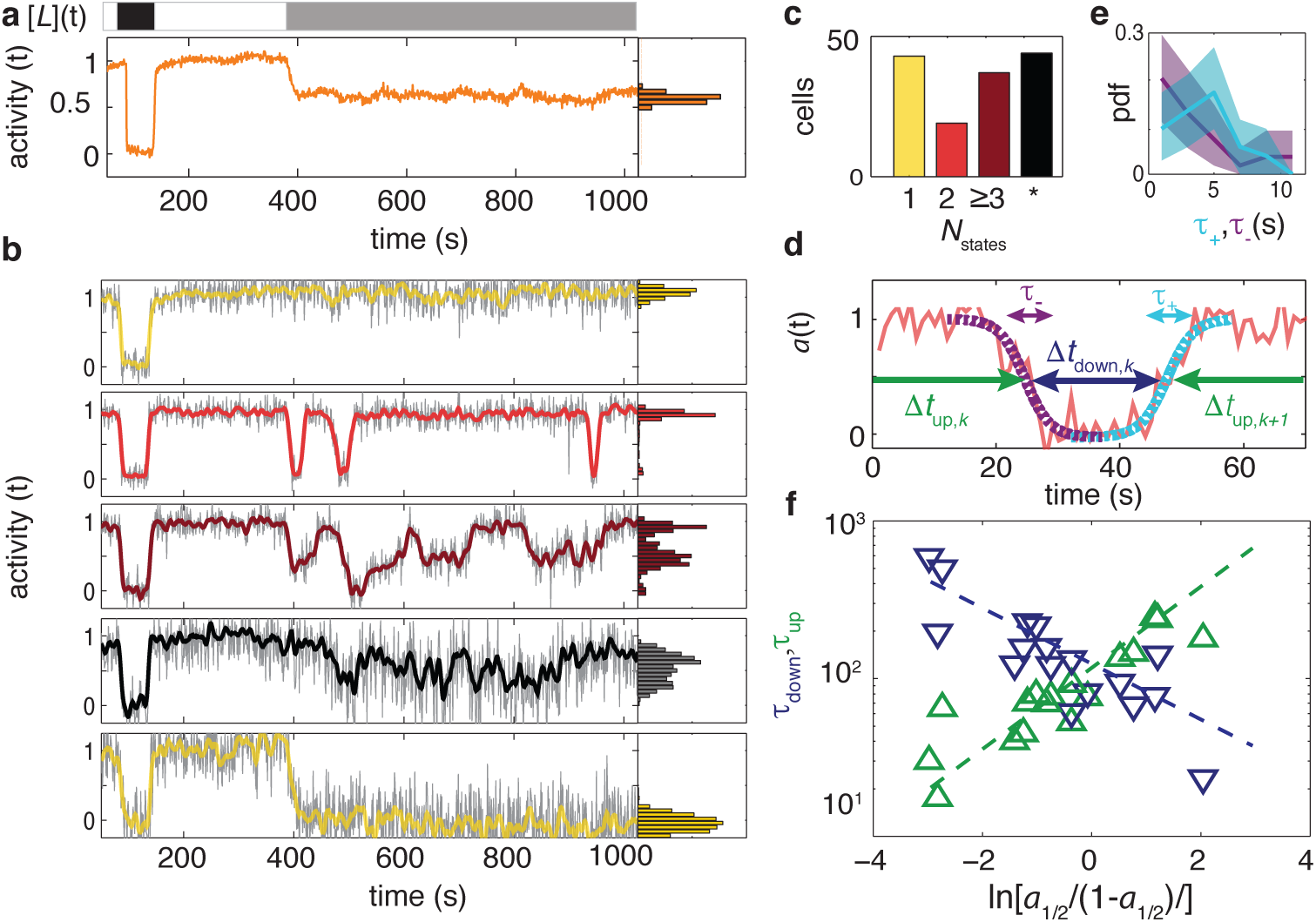
Temporal fluctuations in adaptation-deficient cells expressing Tsr as the sole chemoreceptor (TSS1964). (**a**) (Top) Step response protocol with L-serine. At the start of the experiment, a high (1mM) step response is applied for a rshort time (black). After flushing buffer for several minutes (white) a 20 μM step is applied for ±10 minutes (grey). (Middle) Population-averaged time series of 58 cells responding to the step response protocol. (**b**) Selected single-cell time series of the population shown in panel (a), each normalized to its baseline activity level before adding the first stimulus. To the unfiltered data (grey) a 7s moving average Filter is applied and superimposed (colored according to categories in panel (c)). All time series and corresponding activity histograms of the same experiment are shown in Figure 5 - **Supplement 1** and **5 - Supplement 2**. (**c**) Categorization of the single-cell responses by the number of stable activity levels during the application of the stimulus near *K* of the population. Many cells show only one stable activity level (yellow), corresponding to either a full or none response. Some cells show two stable states (red) or more (purple). In other cells stable states did not show stable states (black). (d) Analysis on cells showing two-state switching to characterize transition times to and residence times in the stable states. Only two-state switchers are analysed in at least 75% of the activity level changed. Transition times *t_+_* and *t_−_* are determined by fitting a sigmodial function (see main text) to the transient part of the time series. Residence times were defined as the interval between two successive transitions, at 50% activity. (**e**) Histogram of transition times, *τ_+_* (4.2 ± 2.2s, 26 events, cyan) and *τ_−_* (3.5 ± 3.2, 29 events, purple) from in total 10 cells of a single experiment with 1 Hz acquisition frequency. (**f**) Residence times *τ_up_* and *τ_down_* as a function of the average activity level *a_1/2_*/(1 - *a_1/2_*). *a_1/2_* is defined as the fraction of time spend in the high activity state. The crossover point, obtained from linear fits of *a*_1_/_2_/(1 – *a*_1/2_) to log(τ), is at 110 ± 10 and the slopes are *γ_up_* = 0.4 ± 0.1 and *τ_down_* −0.6 ± 0.1 =. All fit results are ± the standard error from the fit. In total 17 cells are analyzed from 3 independent experiments (one at 1 Hz sampling, two at 0.2 Hz sampling). **Figure 5 - Supplement 1** All single-cell FRET time series from a single representative experiment. **Figure 5 - Supplement 2** Histograms of activity during attractant stimulus for all single cells from a representative single experiment.1092 Supplemental Information

We further noted that within this subset of cells with large temporal fluctuations, a large fraction (54/98) demonstrated fluctuations that resemble rapid step-like transitions between discrete levels of relatively stable activity that could be identified as peaks in the distribution of activity values across time (Fig. 5b, marginal histograms). Among these “stepper” cells, the majority (37/54) appeared to transition between 3 or more discrete activity levels (Fig. 5b, brown curve), whereas the remaining sizable minority of steppers (17/54) demonstrated binary switching between two discrete levels corresponding to to the maximum (*a* ≈ 1) and minimum (*a* ≈ 0) receptor-kinase activity states (Fig. 5b, red curve). The remaining fraction of cells (44/98) demonstrated fluctuations that were also often large but in which discrete levels could not be unambiguously assigned (Fig. 5b, black curve). The numbers of cells corresponding to each of the categories described above are summarized in Fig. 5c.

The observation of cells that demonstrate spontaneous two-level switching is particularly surprising, given the large number of molecules involved in receptor-kinase signaling. The expression level of each protein component of the chemoreceptor-CheW-CheA signaling complex in our background strain (RP437) and growth medium (TB) has been estimated (by quantitative Western Blots) to be of order 10^4^ copies/cell (***Li and Hazelbauer, 2004***). Considering that the core unit of signaling has a stoichiometric composition of receptor:W:A = 12:2:2 (monomers) (***Li and Hazelbauer, 2011***), the number of core units is likely limited by the number of receptors, leading to an estimate 10^4^/12 ∼ 10^3^ core units for a typical wildtype cell. This estimate does not apply directly to the experiments of Fig. 5 because receptors are expressed from a plasmid in a strain deleted for all receptors. But the FRET response amplitudes of these cells were similar to those of cells with a wildtype complement of receptors, and we thus expect the number of active core units per cell in the experiments of Fig. 5 to be similar to or greater than that in wildtype cells.

We analyzed further the temporal statistics of the discrete transitions in the subset of cells exhibiting two-level switching (Fig. 5d-f). We first quantified the duration of time over which such transitions in activity occur by fitting segments of the activity time series over which these switches occured (Fig. 5d) with a symmetrized exponential decay function (see **Materials and Methods**) to obtain switch durations *τ*_+_ and *τ_-_* for upward and downward transitions, respectively. The fitted values for *τ*_+_ and *τ_-_* correspond to the duration over which the activity trajectory traverses a fraction 1 - *e^-1^* of the transition’s full extent, and were found to be similar between switches in both directions: 〈*τ*_+_〉 ± *σ_τ_*_+_ = 4.2 ± 2.2 s and 〈*τ_-_*〉 ± *σ_τ__* = 3.5 ± 3.2 s (Fig. 5e).

We then considered the duration of time between switching events. We defined Δ*t_up,k_* and Δ*t_down,k_* as the duration of the k-th time interval between transitions with high- and low-activities, respectively (Fig. 5d), and computed the average over all *k* of Δ*t_up_/_down,k_* for each individual cell to estimate its residence timescales τ*_up/down_* for states of high/low activity, respectively. From each cell’s set of intervals *{Δ*t_up/down,k_*}* we also computed a parameter *a*_1/2_, defined as the fraction of time the cell spent in the high activity level, as a measure of its time-averaged activity during the sub-saturating (20μM) L-serine stimulus that yielded a population-averaged response 〈*a*〉 ≈ 1/2 (see **Materials and Methods**).

We found that the logarithms of the residence times scale approximately linearly with ln[*a*_1/2_/(1 - *a*_1/2_)] (Fig. 5f). The latter can be considered a free-energy difference (-Δ*G*) = *G_down_ - G_up_* between the inactive and active states of an equilibrium two-state switching process in which the time-averaged activity *a_1/2_* is given by the probability of being in the active state, *a_1/2_* = *p*(active) = [1 + *e^ΔG^*]^−1^. The residence time in each state can then be described by an Arrhenius-type relation with attempt rate for barrier crossing *τ_ν_* and the height of the energy barrier, 
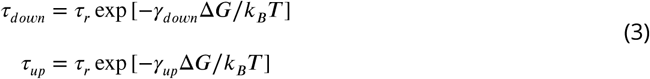
in which the constants *γ_down_* and *γ_up_* describe how the barrier heights of the down and up states depend on the free-energy difference *ΔG* = ln[(1−*a_1/2_)/a_1/2_*]. We find *γ_down_* = − 0.4±0.1, *γ_up_* = 0.6±0.1, and the attempt timescale *τ_r_*, defined here as equivalent to *τ_up_* = *τ_down_* when *a_1/2_* = 1/2, was found to be 110 ± 10 s.

In summary, these data demonstrate the existence of a signaling noise source that is independent of the adaptation enzymes CheR/CheB. The fluctuations they generate can be very strong in cells expressing Tsr as the sole chemoreceptor, leading to two-level switching in a subset of cells. The latter observation suggests that cooperativity among signaling units in chemoreceptor arrays can reach extremely high values, with up to ∼ 10^3^ units switching in a cooperative fashion. The temporal statistics of these two-level switches are consistent with a barrier-crossing model in which the residence time of both states depend on the activity bias *a*_1/2_.

## Discussion

The single-cell FRET measurements described here allowed us to quantify variability in a variety of signaling parameters of the bacterial chemotaxis system, both across cells in a population and within individual cells over time. By imaging many (up to ∼ 100) cells simultaneously, we are able to collect single-cell statistics at high throughput to build up single-cell statistics. Although single-cell experiments have a long history in studies of bacterial chemotaxis (***Berg and Brown, 1972; Spudich and Koshland, 1976; Block et al., 1982; Korobkova et al., 2004; Dufour et al., 2016***),nearly all examples to date have relied on measurements of flagellar motor output (in either tethered or swimming cells). A major advantage of the FRET approach is that it provides a direct measurement of intracelluar signaling that bypasses the noisy behavior of the flagellar motor (a stochastic two-state switch), thereby enabling accurate and efficient determination of signaling parameters.

### From gene-expression noise to network-level diversity

A key feature of bacterial chemotaxis as an experimental system is that one can study *in vivo* signaling and behavior in a manner that is decoupled from gene expression and growth. Being an entirely protein-based signaling network, chemotaxis signaling responses do not require changes in gene expression, and the relatively short timescales of signaling reactions (subsecond to minutes) are well separated from those of changes in protein counts due to gene expression noise (minutes to hours). The ensemble of single-cell FRET time series measured in each of our experiments thus provide a snapshot of cell-to-cell variability due to stochastic gene expression in a variety of signaling parameters.

Our data revealed high variability in important signaling parameters connected to the adaptation system (Fig. 1). In the case of the variability in recovery times (*CV*=0.20), this is likely due to variability in the CheR/receptor ratio from cell to cell. What consequences might such variability have on chemotactic behavior? A recent theoretical study has established that long (short) adaptation times are better suited for maximizing chemotactic migration rates in shallow (steep) gradients (***Frankel et al., 2014***). Thus, variability in adaptation times could partition the population into cells that will be more efficient in running up steep gradients, while others are specialists in climbing shallow ones. Interestingly, it was also found that optimal performance at each gradient involves tuning not only the adaptation time, but also other parameters such as swimming speed or tumble bias, leading to a selective pressure not only for the distribution of individual parameters, but also correlations among them (***Frankel et al., 2014; Waite et al., 2016***). Whether such correlated variation exists among signaling parameters would be a fruitful avenue for future single-cell FRET studies.

In the ligand response of the network, we observed large cell-to-cell variability in the sensitivity (1/K) and steepness (*H*) of dose-response relations, for cells with a wildtype receptor population (Fig. 2). Using a mixed-species MWC model (***Mello and Tu, 2005***), we were able to estimate the Tar/Tsr ratio in single cells, which spans a broad range from nearly zero to more than two. This strong variability in the receptor-cluster composition has the potential to dramatically impact behavior. In their natural habitats, cells likely experience a variety chemoeffector gradients simultaneously, each associated with an unknown fitness payoff for chemotactic pursuit. Generating diversity in the chemoreceptor ratio, which has been shown to determine which gradient to climb when challenged with such conflicting possibilities (***Kalinin et al., 2010***), could allow the isogenic population to hedge its bets to maximize net fitness gains. The Tar/Tsr ratio has also been shown to play an important role in setting the preferred temperature for thermotaxis (***Salman and Libchaber, 2007***; ***Yoney and Salman, 2015***). Variability in Tar/Tsr would allow diversification of the preferred temperature across cells in the population, which will promote spreading of bacteria in environments with temperature gradients. Finally, when chemotactic bacteria colonize an initially nutrient-rich environment, they are known to successively exploit resources by emitting multiple traveling waves of chemotactic bacteria, each of which consumes and chases by chemotaxis a different nutrient component outward from the colony origin (***Adler, 1966***). Our observation that the population diversity in receptor ratios, and hence chemotactic preference, varies concomitantly with population growth could provide a means to tune the population fractions that engage in such excursions into virgin territory, and those that remain for subsequent exploitation of remaining resources. Thus, the diversity in ligand response and preference generated by variability in the Tar/Tsr ratio could have nontrivial consequences in a variety of behavioral contexts encountered by isogenic chemotactic (and thermotactic) populations.

### Suppression of gene expression noise by CheB phosphorylation feedback

The role of phosphorylation feedback has been a long standing open question in the field of bacterial chemotaxis signaling, ever since its presumed role in providing precise adaptation was decisively ruled out by ***Alon et al. (1999***). In the ensuing years, a diverse set of hypotheses have been proposed to explain its purpose. Apart from precise adaptation, CheB phosphorylation has been suggested as possibly responsible for the non-linear response of CheB activity to changes in CheA kinase activity (***Shimizu et al., 2010; Clausznitzer et al., 2010***), ligand sensitivity of wild-type cells (***Barkai et al., 2001***), and has been implicated theoretically as a possible mechanism to buffer gene-expression noise to suppress detrimental variability in the steady-state kinase activity (***Kollmann et al., 2005***). Here, we tested the latter hypothesis, by severing the phosphorylation feedback loop as a possible noise-reduction mechanism. Our single-cell FRET data revealed that, not only does CheB phosphorylation feedback strongly attenuate the magnitude of variability in the steady-state kinase activity *a*_0_, it also qualitatively changes the shape of the distribution *P*(*a*_0_) across cells to convert an otherwise bimodal distribution into a unimodal one (Fig 3d). The highly polarized bimodal distribution of steady-state activities in CheB phosphorylation mutants are likely detrimental, as they could drive *a*_0_ of a large fraction of the population too far from the flagellar motor’s steep response threshold (***Cluzel et al., 2000; Yuan and Berg, 2013***) to effectively control swimming. The observation of a bimodal *P*(*a*_0_) in the absence of phosphorylation feedback is consistent with a previous modeling study by ***Emonet and Cluzel (2008)*** in that the parameters of the CheR- and CheB-catalyzed covalent modification cycle appear to satisfy conditions for zero-order ultrasensitivity (***Goldbeter and Koshland, 1981***), which has been hypothesized to provide a source of slow and large amplitude temporal fluctuations with possible benefits for chemotaxis in certain environments (see below). The fact that CheB phosphorylation seems to strongly attenuate the steepness of the ultrasensitive relationship *a*_0_ = *f* ([_R]/[B]) between kinase activity and the [R]/[B] ratio suggests that zero-order might not suffice to explain the large amplitude temporal fluctuations we observed in wildtype cells (see below).

### Diversity in temporal noise: bet-hedging across exploration and exploitation strategies

In addition to cell-to-cell variability in signaling parameters, single-cell FRET allowed us to resolve temporal fluctuations in signaling about the steady-state output within individual cells. In wild-type cells, we found that the steady-state activity fluctuates slowly (Fig. 4,correlation time *τ* ≈ 10_s_) with a large amplitude (η = *σ_a_/*〈*a*〉 ≈ 40%), but this amplitude also varies significantly from cell to cell (*CV* ≈ 0.6). Fluctuations on this timescale were absent in CheRB-cells defective in receptor methylation/demethylation, indicating that these fluctuations are generated by stochastic processes in the activity of the adaptation enzymes CheR and CheB. Whereas the fluctuation correlation time τ in our FRET experiments was in close agreement with those from previously reported flagellar motor switching experiments (***Korobkova et al., 2004; Park et al., 2010***), the fluctuation amplitude 〈*η*〉 ≈ 40% was surprisingly large. Theoretical analysis of the motor-based noise measurements (***Tu and Grinstein, 2005***) had predicted a more modest noise level of intracellular noise, at 10-20% of the mean. The discrepancy is likely due, at least in part, to the recently discovered adaptation at the level of the flagellar motor (***Yuan et al., 2012***), which must effectively act as a highpass filter that attenuates frequencies near or below a cutoff frequency determined by its own characteristic timescale for adaptation. The fluctuation amplitude*η* was also much greater than previous estimates from pathway-based models. A possible explanation is the amplification of noise by receptor cluster cooperativiy, which our results show to be much higher compared to previous estimates based on population-averaged measurements. Another possibility is an additional source of noise. In cells with a pure cluster composition we clearly demonstrate this possibility. Investigating how both noise sources contribute in the case of mixed receptor clusters (e.g. CheRB-cells) and the amplification magnitude (dose response curves in CheRB+ cells) should be an interesting direction for future study.

The temporal noise we observed could have profound implications for *E. coli’s* random-walk motility strategy, because slow fluctuations in the intracellular signal can enhance the likelihood of long run events and stretch the tail of the run-length distribution to yield power-law-like switching time distributions over a range of time scales (***Korobkova et al., 2004; Tu and Grinstein, 2005***).Such non-exponential statistics are known to yield superior foraging performance in environments where resource distribution is sparse (***Viswanathan et al., 1999***), and temporal fluctuations in run-tumble behavior has also been shown theoretically to enhance climbing of shallow gradients by generating runs that are long enough to integrate over the faint gradient a detectable difference in ligand input (***Flores et al., 2012; Sneddon and Emonet, 2012***). Hence, the noise generated by the adaptation system can be advantageous in resource-poor environments (*deserts*) in which efficient exploration of space for sparsely distributed sources (*oases*) is of utmost importance. By contrast, strong temporal noise clearly degrades response fidelity in rich environments where the gradient signal is strong enough for detection with short runs, and might also complicate coordination of cells in collective behaviors such as the aforementioned traveling-wave exploitation of nutrients. Our finding that the noise amplitude varies strongly from cell to cell thus suggests that isogenic populations might be hedging their bets by partitioning themselves between specialists for local exploitation of identified resource patches and those for long-range exploration in search for new ones.

### Giant fluctuations and digital switching in adaptation deficient cells

We found the most dramatic temporal fluctuations in adaptation-deficient (CheRB-) cells expressing Tsr as the sole chemoreceptor species (Fig. 5). When brought close to their dose-response transition point (K) by attractant stimulation, these cells demonstrated strong temporal fluctuations, revealing that there exist sources of signal fluctuations that are independent of CheR and CheB activity. The origin of these adaptation-independent fluctuations remain unknown, but in broad terms, one can envisage that they are due to either intrinsic sources (i.e. fluctuations arising within the components of the receptor-kinase complex), extrinsic sources (i.e. fluctuations in other cellular processes / environmental variables), or both. Possible to intrinsic sources include coupled fluctuations in protein conformations (***Duke and Bray, 1999; Shimizu et al., 2003; Mello et al., 2004; Skoge et al., 2011***), the slow-timescale changes in receptor “packing” that have been observed by fluorescence anisotropy measurements(***Frank and Vaknin, 2013; Vaknin, 2014***), and the stochastic assembly dynamics of receptor clusters (***Greenfield et al., 2009***). Possible extrinsic sources include fluctuations in metabolism, membrane potential, or active transport/consumption of ligand. Many of these possibilities could be tested by experiments of the type presented here with appropriate mutant strains and environmental controls, and present promising directions for future research.

The adaptation-independent fluctuations we observed were not only large in amplitude but often (though not always) took the form of discrete steps in activity, in some cases between only two levels. Two-state descriptions of receptor signaling are a common feature of nearly all mechanistic models of bacterial chemotaxis signaling addressing both cooperativity (***Duke and Bray, 1999; Shimizu et al., 2003; Mello et al., 2004; Mello and Tu, 2005; Keymer et al., 2006***) and adaptation (***Asakura and Honda, 1984; Barkai and Leibler, 1997; Morton Firth et al., 1999; Endres and Wingreen, 2006; Tu et al., 2008***), yet direct evidence for two-state switching by receptor-kinase complexes has been lacking. Although as noted above, it is yet possible that the two-level switching we observed (Fig. 5a,b) is due to extrinsic noise sources (e.g. metabolism or transport), the temporal statistics we observed (Fig. 5d-f) are compatible with a simple model in which two stable signaling states are separated by an energy barrier sensitive to both environmental stimuli and internal cell variables.

Regarding cells that exhibited step-like transitions among more than two stable states, a plausible interpretation is that the underlying transitions are actually two-level, but the receptor-kinase population is partitioned into two or more disjoint signaling arrays, and that the observed FRET signals represent the sum of the stochastic two-level activities of these independently switching arrays. For cells in which there is sufficient spatial separation between chemoreceptor clusters, it could be possible to test this hypothesis by conducting on the same set of cells both FRET measurements and receptor-cluster imaging to relate the number of distinct step sizes to the number of observed clusters.

If the stochastic two-level switching we observed is indeed due to intrinsic sources of noise, it would strongly suggest (as discussed in Results) that at least many hundreds, if not thousands of receptor-kinase units are switching in a cooperative fashion. The rather long timescale associated with intervals between switches (≈ 10^2^ s) is indicative of their large size, and is also clearly distinct from the methylation-dependent fluctuation timescale (≈ 10 s) observed in CheRB+ cells. The switching duration (≈4 s), is also much slower than the sub-second response to attractant stimuli (***Segall et al., 1982; Sourjik and Berg, 2002a***). Interestingly, these timescales (∼ 10^2^ s and ∼ 10^0^ s for inter-switch intervals and switch durations, respectively) are approximately 10^2^-fold longer than those measured by ***Bai et al. (2010***) for the two-state switching of the flagellar motor (∼ 10^0^ s and ∼ 10^-2^ s for inter-switch intervals and switch durations, respectively), whose C-ring is composed of ∼ 10^1^ allosteric units, approximately 10^2^-fold less than the ∼ 10^3^ units we estimate for the number of receptor-kinase units per cell in our experiments. The study of ***Bai et al. (2010***) demonstrated impressive agreement between those temporal statistics and predictions of a “conformational spread” model (***Duke et al., 2001; Bray and Duke, 2004***), an adaptation of the equilibrium Ising model for ferromagnetism (***Ising, 1925***). It would be interesting to investigate whether similar Ising-type models for the receptor lattice (***Duke and Bray, 1999***; ***Shimizu et al., 2003***; ***Mello et al., 2004***; ***Skoge et al., 2006***) can explain the timescales observed in our receptor-kinase switching data.

Although our results indicate that, at least to a first approximation, receptor-kinase switching can be treated as a thermally driven barrier-crossing process, we note that our data do not rule out the possibility of non-equilibrium switching mechanisms (***Tu, 2008***). Indeed, despite the success of equilibrium models in closely matching a wealth of data on the flagellar motor switch (***Bai et al., 2010***), recent experiments have revealed new evidence that switching of the motor C-ring likely includes also an active component, effectively utilizing part of the energy dissipated in motor torque generation to enhance sensitivity (***Wang et al., 2017***). Like the motor C-ring switch, whose state is only observable when coupled to rotation driven by dissipative proton conduction, essentially all experimental methods available to study receptor-kinase signaling involve coupling to a dissipative process (ATP hydrolysis by CheA) for readout. The experimental access to receptor-kinase temporal statistics afforded by single-cell FRET holds promise to help discriminate possible equilibrium and non-equilibrium mechanisms for signal processing within this remarkable protein circuit.

### Concluding remarks

We described a new single-cell FRET technique capable of resolving intracellular signaling dynamics in live bacteria over extended times. Our results highlight how a protein-based signaling network can either generate or attenuate variability, by amplifying or filtering molecular noise of different molecular origins. Gene expression noise is harnessed, on the one hand, to generate diversity in the ligand response of isogenic populations, or attenuated, on the other the hand, in the control of steady-state signal output. In addition, we showed that signaling noise generated at the level of interacting gene products can have a profound impact. Stochastic protein-protein interactions within the signaling network, as well as other "extrinsic" fluctuations, can be amplified by the signaling network to generate strong temporal temporal fluctuations in the network activity.

## Materials and Methods

### Strains and Plasmids

All strains used are descendants of *E. coli* K-12 HCB33 (RP437). Growth conditions were kept uniform by transforming all strains with two plasmids. All strains and plasmids are shown in Tables 1 and 2.

**Table 1.**
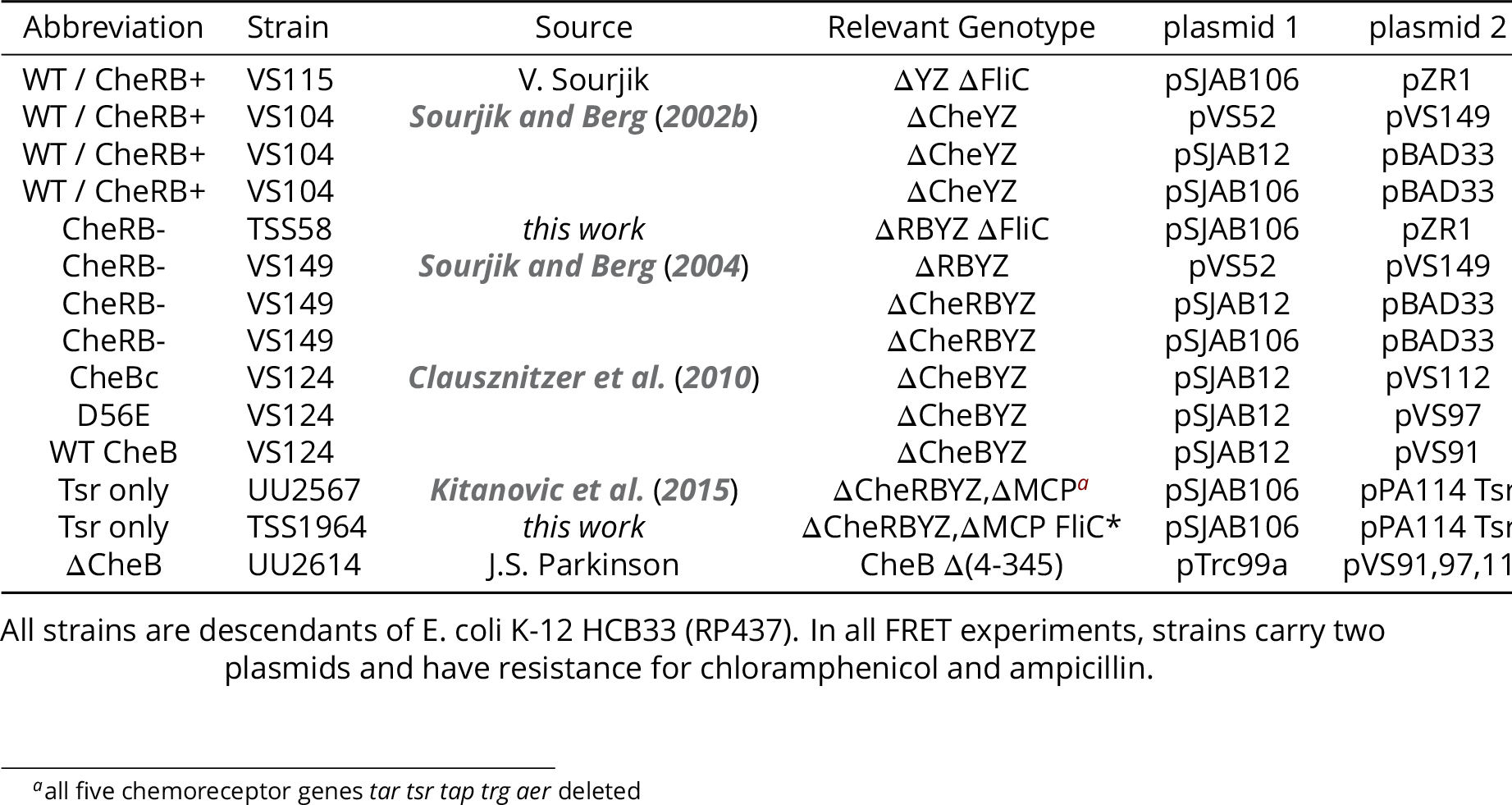
Strains used in this study.

**Table 2.** Plasmids used in this study.

| Name | Product | System | Ind | Res | Source |
| --- | --- | --- | --- | --- | --- |
| pVS52 | CheZ-5G-YFP | pBAD33 | ara | cam | *Sourjik and Berg (2002b)* |
| pVS149 | CheY-5G-mRFP1 | pTrc99a | IPTG | amp | *Sourjik and Berg (2002b)* |
| pSjAB12 | CheZ-5G-YFP / CheY-5G-mRFP1 | PTrc99a | IPTG | amp | This work |
| pSjAB106 | CheZ-5G-YFP <sup>a</sup> / CheY-5G-mRFP1 | PTrc99a | IPTG | amp | This work |
| pVS91 | CheB <sup>b</sup> | pTrc99a | ara | cam | *Liberman et al. (2004)* |
| pVS97 | CheB-D56E <sup>c</sup> | pBAD33 | ara | cam | *Clausznitzner et al. (2010)* |
| pVS112 | CheBc <sup>d</sup> | pBAD33 | ara | cam | V. Sourjik |
| pSjAB 122 | CheBc-GS4G-mVenus | pBAD33 | ara | cam | This work |
| pSjAB 123 | CheB(D56E)-GS4G-mVenus | pBAD33 | ara | cam | This work |
| pSjAB 124 | CheB-GS4G-mVenus | pBAD33 | ara | cam | This work |
| pZR 1 | FliC* <sup>e</sup> | pKG116 | NaSal | cam | This work |
| pPA114 Tsr | Tsr | pPA114 | NaSal | cam | *Ames et al. (2002)* |

<sup>a</sup>Contains a A206K mutation to enforce monomerity.
<sup>b</sup>expresses WT CheB
<sup>c</sup>carries a point mutation D56E in CheB
<sup>d</sup>expresses only residues 147-349 of CheB, preceded by a start codon (Met)
<sup>e</sup>expresses sticky FliC element *Scharf et al. (1998)*

The FRET acceptor-donor pair (CheY-mRFP and CheZ-YFP) is expressed in tandem from a IPTG inducible pTrc99A plasmid, pSJAB12 or pSJAB106, with respective induction levels of 100 and 50 μM IPTG. The differences between pSJAB12 and pSJAB106 are i) the presence of a noncoding spacer in pSJAB106 to modify the ribosome binding site of CheZ ***Salis et al. (2009)***, such that CheZ is expressed approximately 3 fold less, and ii) a A206K mutation in YFP to enforce monomerity. We also used pVS52 (CheZ-YFP) and pVS149 (CheY-mRFP1) to express the fusions from separate plasmids with induction levels of 50 uM IPTG and 0.01 % arabinose, respectively. We transformed the FRET plasmids in adaptation-proficient strain (VS104) to yield CheRB+ and adaptation-(VS149) to get CheRB-. For attachment with sticky flagella from pZR1 we used the equivalent strains in *fliC* background (VS115 and TSS58).

Experiments with Tsr as the sole chemoreceptor were performed in UU2567 orTSS1964, in which the native FliC gene is changed to sticky FliC (FliC*). Tsr is expressed from pPA114Tsr, a pKG116 derivative, at with an induction of 0.6 μM NaSal.

For the experiments with the CheB mutants, pSJAB12 was transformed into VS124 together with plasmids expressing CheB^WT^, CheB^B56E^ and truncated mutant CheB_c_ (plasmids pVS91, pVS97 and pVS112, respectively, with induction levels of 1.5E-4, 6E-4 and 3E-4 % arabinose.

All strains are descendants of E. coli K-12 HCB33 (RP437). In all FRET experiments, strains carry two plasmids and have resistance for chloramphenicol and ampicillin.

### FRET Microscopy

Föster Resonance Energy Transfer [FRET] microscopy was performed as previously reported (***Sourjijk et al., 2007; Vaknin and Berg, 2004***). Cells were grown to OD=0.45-0.5 in Tryptone Broth (TB) medium from a saturated overnight culture in TB, both with 100 μg/mL ampicillin and 34 μg/mL chloramphenicol and appropriate inducers in the day culture. For the FRET experiments we used Motility Media (MotM) (***Shimizu et al., 2006***), in which cells do not grow and protein expression is absent. Cells were washed in 50 mL MotM, and then stored 0.5-6 h before experiment. In the dose-response curve experiments and the temporal fluctuation measurements, cells were stored up to three hours at room temperature to allow for further red fluorescence maturation. A biological replicate or independent FRET experiment was defined as a measurement from separately grown cultures, each grown on a separate day.

Cells were attached by expressing sticky FliC elements from a pKG116 plasmid, induced with 2μM Sodium Salicylate (NaSal), or with Poly-L-Lysine (Sigma), or with anti-FliC antibodies column purified (Using Protein A sepharose beads, Amersham Biosciences) from rabbit blood serum and pre-absorbed to FliC-cells (HCB137, gifts from Howard Berg). We found FRET experiments with sticky FliC to have the highest signal-to-noise ratio.

Fluorescent images of the cells were obtained with a magnification of 40-100x (Nikon instruments). For excitation of YFP, we either used 514 nm laser excitation set to 30 mW for 2 ms or an LED system (CoolLED, UK) with an approximate exposure time of 40 ms to approximate the same illumination intensity per frame. The sample was illuminated stroboscopically with a frequency between 1 and 0.2 Hz. RPF excitation was performed by 2ms exposure of 60 mW 568 nm laser or equivalent with LED to measure acceptor levels independently from FRET.

Excitation light was sent through a 519 nm dichroic mirror (Semrock, USA). Epifluorescent emission was led into an Optosplit (Cairn Research, UK) with a second dichroic mirror 580 nm and two emission filters (527/42 nm and 641/75 nm, Semrock, USA) to project the RFP and YFP emission side by side on an EM-CCD (Princeton Instruments, USA) with multiplication gain 100. All devices were controlled through custom-written software.

### Image processing

Images were loaded and analyzed by means of in-house written scripts in MATLAB and Python. Images were corrected for inhomogeneous illumination. Single cells were selected by image segmentation on the donor emission with appropriate filter steps to remove clusters of cells or cells improperly attached to the coverslip. At the position of each cell a rectangular ROI is defined in which all fluorescence intensity is integrated. For experiments in which the concentration of donor molecules may influence the FRET signal, the experiments on the CheB mutants, segmentation was done separately for each frame to determine the cell shape and then linking these segmented images with a tracking algorithm (***Crocker and Grien, 1996***), afterwards, fluorescence intensities are normalized for the cell size (mask surface area) in segmentation and cells with low acceptor intensities were excluded from the analysis. The ROI for the donor intensity were subsequently used to obtain the acceptor intensity per cell, both in photon-count per pixel. Fluorescence intensities were corrected for bleaching by fitting a linear, single exponential or double exponential function to the fluorescence decay, separately for both donor and acceptor. Cells in which the intensity decay cannot accurately be corrected were excluded from the analysis.

### FRET analysis

The FRET signal is calculated from fluorescent time series. We observe changes in the ratio *R* = *A/D*,in which A and D are the fluorescence intensities of the acceptor and donor. In previous population-averaged FRET experiments the FRET per donor molecule (Δ*D/D_0_*) is calculated as (***Sourjik and Berg, 2002b; Sourjijk et al., 2007***):

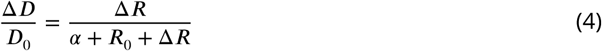
in which *R_0_* is the ratio in absence of FRET, *α* = |Δ*A*/Δ*D*| is a constant that depends on the experimental system (in our case *α* = 0.30) and the change in ratio as a result of energy transfer Δ*R*. Δ*R* and *R_0_* are obtained through observing the ratio just after adding and removing saturated attractant stimuli. This expression is convenient for population FRET since is invariant to attachment densities of a population. However, in single-cell FRET this expression may generate additional variability in FRET due to variable donor levels from cell to cell. Hence it is more convenient to define the FRET levels in terms of the absolute change in donor level ΔD, since this reflects the number of resonance energy transfer pairs

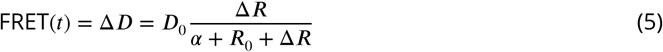

Since FRET occurs only when CheY-P and CheZ interact, the FRET level is proportional to the concentration of complex [Yp-Z]. If we assume the CheY-P dephosphorylation by CheZ follows Michaelis-Menten kinetics we can describe the [Yp-Z] concentration in terms of the activity of the kinase CheA. For this, we assume the system is in steady-state for timescales much larger than CheY phosphorylation-dephosphorylation cycle (≈ 100 ms). In that case, the destruction rate should equal the rate of CheA phosphorylation and hence the FRET signal is proportional to the activity per kinase *a* and the amount of CheA in the kinase-receptor complex (***Sourjik and Berg, 2002b;*** Oleksiuk et al., 2011):

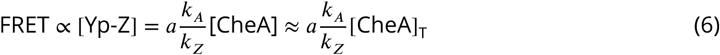

This last step is only valid if we further assume CheA autophosphorylation being the rate-limiting step. This is the case only if sufficient amounts of CheZ and CheY present in the cell. We have found that the FRET level initially increases with donor (CheZ) levels, but then saturates and remains constant for CheY and CheZ (see Fig. 3 - Supplement 2).

In many cases the most relevant parameter is the normalized FRET response. The FRET level reaches maximum if all kinases are active (a ≈ 1). In case of CheRB+ cells, this is the case when removing a saturating amount of attractant after adaptation (***Sourjik and Berg, 2002b***). For CheRB-cells the baseline activity is (***Sourjik and Berg, 2002b; Shimizu et al., 2010***) close to 1. Hence the normalized FRET FRET(*t*)/FRET_max_ represents the activity per kinase *a*(*t*) and is the relevant parameter for many quantitative models for chemoreceptor activity (***Tu, 2013***).

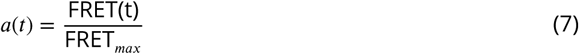
and from *a*(*t*) the steady-state activity *a_0_* can be determined by averaging *a*(*t*) over baseline values before adding attractant stimuli.

### Power Spectral Density Estimates

From FRET time series of length *T* and acquisition frequency *f* we calculated Power Spectral Density [PSD] estimates as

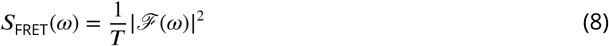
in which *ℱ*(*ω*) is the discrete-time Fourier transform. We only consider positive frequencies and multiply by two to conserve power.

To study the influence of experimental noise and the effect of estimating *τ* and *c* from a finite time window, we generated O-U time series using the update formula (***Gillespie, 1996***)

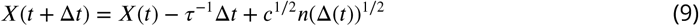
in which *n* denotes a sample value from a normal variable. To the generated time series Gaussian white noise was added to simulate experimental noise. The experimental noise amplitude was obtained from the average power at high frequencies.

### Two-state switching analysis

We obtained switching durations by fitting the function

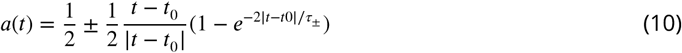
to the normalized FRET time series in a 30-second time window, approximately ±15 s from t_0_. The residence times Δ*t_up,i,k_* and Δ*t_down,i,k_* of event *k* in cell *i* were defined by the time between transitions or the beginning/end of the 20 *μM* stimulus time window. The steady-state activity during activity was then calculated as 
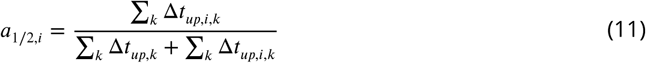
and for the residence times we take the mean over *k* to get *τ_down_* and *τ_up_*. If we treat the system as an equilibrium process we can use the Arrhenius equations that describe the residence times as a function of the distance to the energy barrier

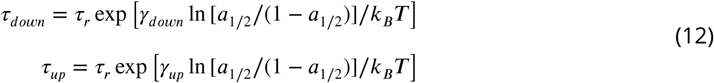
in which *γ_down_* and *γ_up_* are constants corresponding to the slopes of ln *τ_down_* and ln *τ_up_* against ln [*a*_1/2_/(1 – *a*_1/2_)], respectively. The At parameters and standard error are obtained with the robustfit function in Matlab (statistics toolbox).

### Dose Response Curve Analysis

Normalized FRET responses to different levels of ligand are fit to a hill curve of the form 
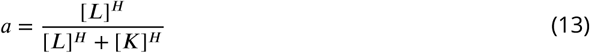

This can be connected to an MWC-type model (**Monod et al., 1965)** of receptor cluster activity (***Shimizu et al., 2010***) in the regime *K_I_* << [L] << *K_A_*, resulting in the correspondence key

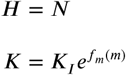
which relates the Hill slope directly to the cluster size N, and sensitivity *K* to the methylation energy of the receptor. We plot *K* on a logarithmic scale to scale linearly with energy.

To obtain expression level estimates of different receptor species we use a different MWC model. Following***Mello and Tu (2005)***, we use as an expression for the normalized response of cells to ligand *[L]* serine 
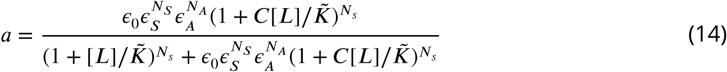
in which *N_A_* is the number of Tar receptors in the cluster and *N_s_* is the number of Tsr receptors Parameters *ε_A_, ε_s_, ε_0_* are the energies corresponding to binding of ligand to Tar, Tsr and the other three receptors and are the same for each cell, like *C* and 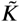 which describe the disassociation constant for the active state as *K_A_* = 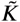/*C*, while *N_A_* and *N_T_* may vary from cell to cell. This yields the minimization problem for all 128 cells 
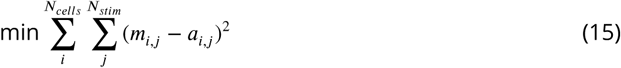
in which *m_ij_* the measured FRET response normalized to the response amplitude of cell *i* to stimulus *L_j_*. This function was minimized using the matlab function fmincon (optimization toolbox). The total number *N_T_* = *N_A_* + *N_s_* is limited to 32. When fitting the model used the energy parameters *ε* from reference ***Mello and Tu (2005)*** where used as initial guess with a maximum of ±5% deviation. This yielded an estimate of *N_A_* and *N_s_* for each cell. Under the assumption that receptor clusters are well-mixed, this yields a Tar/Tsr ratio of *N_A_/N_S_*.

**Table 3.** List of global parameters used for model of Mello and Tu. In these fits, 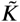 is a free parameter while others are constrained ±5% by published values.

| Parameter | Start Value *Mello and Tu (2007)* | Final Value |
| --- | --- | --- |
| $C$ | 0.314 | 0.29 |
| $\epsilon_0$ | 0.80 | 0.84 |
| $\epsilon_A$ | 1.23 | 1.29 |
| $\epsilon_S$ | 1.54 | 1.61 |
| $\tilde{K}$ | – | 21.2 $\mu\text{M}$ |

### Ultrasensitive adaptation model of phoshorylation feedback

We model the methylation-demethylation cycle of the receptors as a Goldbeter-Koshland cycle (***Goldbeter and Koshland, 1981; Emonet and Cluzel, 2008***). For simplicity, we do not explicitly describe the methylation and demethylation of the receptors explicitly but instead assume that CheR (R) activates the receptor-kinase complex directly (A^*^), and that CheB (B) deactivates it (A).

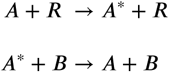

We now assume that these reactions follow Michaelis-Menten kinetics and the total amount of kinase complexes is constant (*A*_T_ = *A\** + *A*). Hence the change in 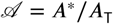 can be described as

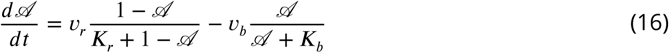
in which *v_r_* and *v_b_* are the rates for the reactions mediated by R and B, respectively. The Michaelis-Menten constants *K_b_* and *K_r_* are in units of *A_T_* and are therefore dimensionless numbers. We are interested in the steady-state level 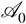 and its dependence on the kinetic parameters in equation 16. This is described by the Goldbeter-Koshland function ***Tyson et al. (2003***), an exact solution to the system in case [R] and [B] are much smaller than [A]_T_.

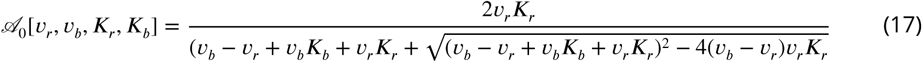

The shape of this curve is sigmodal if the Michaelis-Menten constants *K_r_* and *K_b_* are much smaller than one. For CheB phosphorylation, we assume the phosphorylation rate depends linearly on active CheA and write

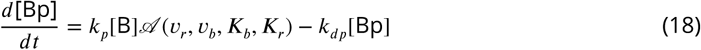

with the corresponding conservation law B_*T*_=B_*P*_+B. For the case for wild-type CheB, with phosphrylation feedback, the rates can be described in terms of catalytic rate times the enzyme (subspec concentration

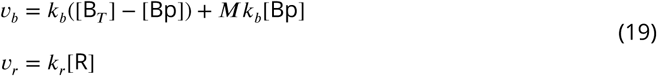

in which *M* stands for the ratio of demethylation rates of unphosphorylated and phosphorylated CheB. The fraction of the phosphorylated CheB, [Bp]/[B]_T_, which is given by *k_dp_*/(*k_p_* + *k_dp_*) thei determines the effective activity of CheB. Equation 18 is solved numerically using Mathematica for [Bp] and the result is substituted in equation 17. In the absence of feedback, the activity can be directly calculated from equation 17 with the rates being simply 
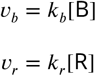

We only need to consider the ratio of rate constants *k_r_* and*k_b_* which determines at which expression ratio [CheR]/[CheB] the activity equals 1/2. We assume *k_r_* = *k_b_* for simplicity, since the shape of the curve from Eq. 17 is not affected, it only shifts the the curve along the horizontal axis. Similarly, we only consider the ratio of phosphorylation and dephosphorylation rates. This leaves the system of equations above only has a few parameters: *K_b,r_;* M; and the ratios *k_r_/b_b_* and *k_p_/k_dp_*, *M*. In table 4, the parameters used for the calculations are listed.

**Table 4.**
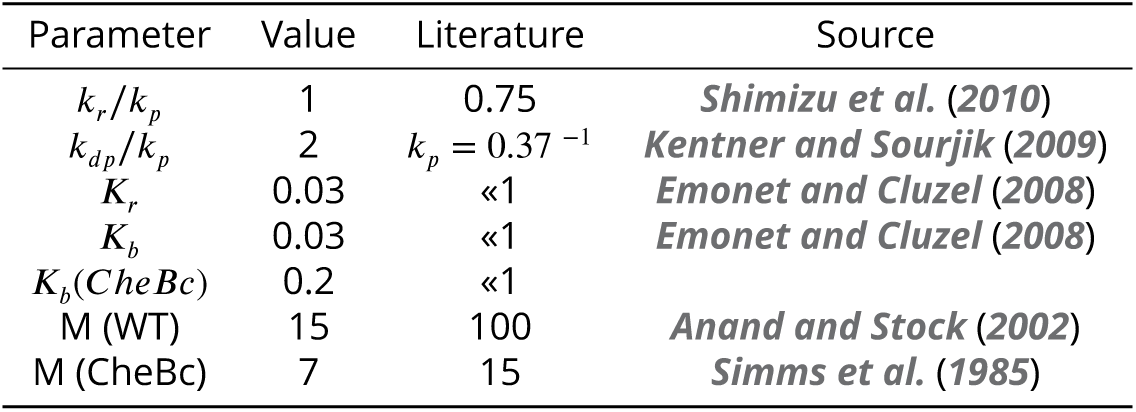
List of parameters used for Goldbeter-Koshland description of CheB phosphorylation feedback.

| Parameter | Value | Literature | Source |
| --- | --- | --- | --- |
| $k_r/k_p$ | 1 | 0.75 | **Shimizu et al. (2010)** |
| $k_{dp}/k_p$ | 2 | $k_p = 0.37^{-1}$ | **Kentner and Sourjik (2009)** |
| $K_r$ | 0.03 | $\ll 1$ | **Emonet and Cluzel (2008)** |
| $K_b$ | 0.03 | $\ll 1$ | **Emonet and Cluzel (2008)** |
| $K_b(CheBc)$ | 0.2 | $\ll 1$ | |
| M (WT) | 15 | 100 | **Anand and Stock (2002)** |
| M (CheBc) | 7 | 15 | **Simms et al. (1985)** |

We first fixed the phosphorylation rates *k_p_* >*2k_dp_*. This means that the steady-state phosphorylated level of CheB [B_p_]/[B_T_] at activity ≈ ^1^/3 is around 15 %. This parameter is not constrained by any direct observation, but it is clear the system benefits from a relatively low fraction of phosphorylation, to be able to up and down regulate the levels effectively upon changes in activity.

Generally, we assume D56E to behave like unphosphorylated CheB. The gain in catalytic rate of activated CheB is estimated to be nearly a 100 fold, but this does not agree with the expression level differences between the different CheB mutants so we made a conservative estimate of 15 (the attenuating effect increases with the gain). CheBc behaves approximately like phosphorylated CheB (albeit with increase of only 7 compared to D56E), qualitatively consistent with measured *in vitro* rates for CheBc and phosphorylated intact CheB ***Anand and Stock (2002)***. The difference between predicted rates and might be due to the fact that the rate experiments were performed ***in vitro.*** Michaelis Menten constants used in the model are lower than 1, but how low is not well constrained by data, and estimations do not take into account the possible attenuating effect of phosphorylation. Our experimental data on the distribution of *a_0_* implies the sigmodial curve is steep in the absence of phosphorylation and hence that *K_b_* and *K_r_* are quite small. The variability in *a_0_* for CheBc is lower than D56E, implying that the curve is less steep and hence we have chosen are *K_r_* which is not quite as low as D56E.

To simulate gene expression noise, we simulated [CheR]/[CheB] log-normal distributions with *σ* = 0.18 for all three strains. The mean of the distribution was chosen to yield an average steady-state network activity (*a*_0_) of 0.4. The resulting distribution of *a_0_* was calculated using the corresponding Goldbeter-Koshland function for each genotype.

## Acknowledgments

We express our gratitude to Simone Boskamp and Zuzana Rychnavska for cell culture and cloning, Marco Kamp for microscopy assistance and Marco Konijnenburg, Luc Blom and Eric Clay for software and electronic support. We also thank Istvan Kleijn, Iwan Vaandrager, Francesca van Tartwijk and Pieter de Haan for help with experiments and William Pontius for useful discussions. We thank Sandy Parkinson and Germán Piñas for many useful discussions and critical reading of the manuscript.

---

**Figure 1 - Supplement 1.**
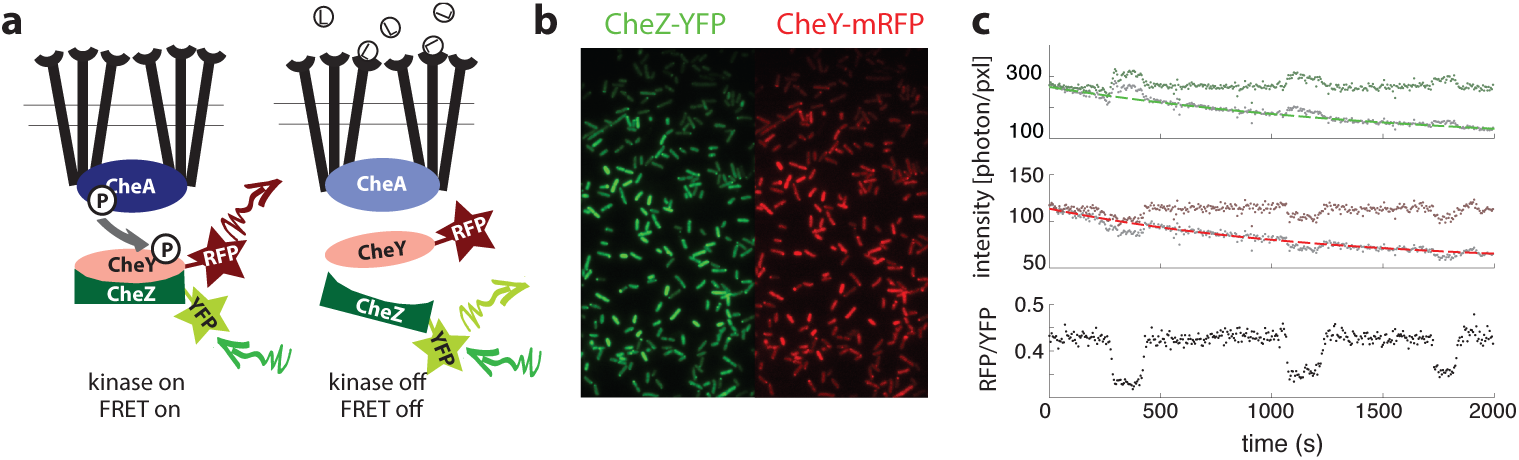
Single-cell FRET assay schematic and workflow. **a**) Schematic of CheY-CheZ FRET assay. In absence of ligand stimuli, receptors are activating CheA. When active, CheA phosphorylates CheY, increasing interaction between CheY and phosphatase CheZ and thereby increasing FRET through the labeled fluorophores. Ligand-receptor binding shuts down the kinase, which ceases phosphate transfer and FRET levels decrease. **b**) False-color images of donor (CheZ-YFP) and acceptor (CheY-mRFP1) fluorescence, channels projected on the same EM-CCD camera chip. **c**) Example time series fluorescence from a single cell (CheRB-,VS149/pVS149/52). In grey raw data is shown, with a Fit to a single exponential function with offset overlaid for donor (top, green) and acceptor channel (middle, red). From the corrected fluorescence intensities the ratio RFP/YFP is calculated (bottom).

**Figure 1 - Supplement 2.**
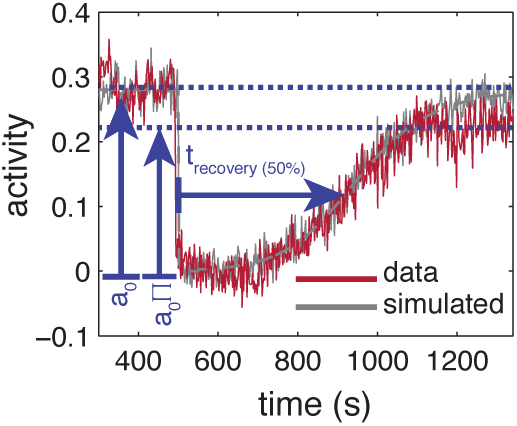
Influence of experimental noise on estimating recovery times. Simulated time series for are calculated by integrating the linearized MWC model with adaptation kinetics (***Tu et al., 2008***) with parameters chosen to closely approximate the population averaged response (grey-dashed line). To the simulated time series of each cell gaussian white noise (*μ* = *0; σ* = 0.15, 55 cells) is added to approximate the noise level of the experiment. Also shown are the baseline and recovery levels (blue dashed lines) of the experimental population.

**Figure 2 - Supplement 1.**
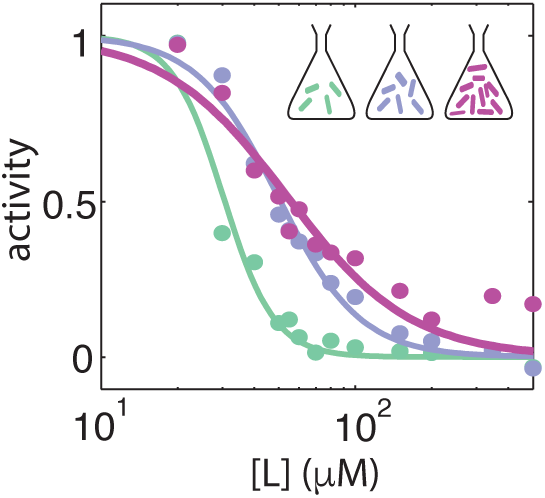
Dose response curves from population-averaged time series at harvesting OD’s 0.31 (green), 0.45 (blue) and 0.59 (purple). The K/H Fit value pairs are respectively 30/4.3, 50 /2.7 and 53/1.8.

**Figure 2 - Supplement 2.**
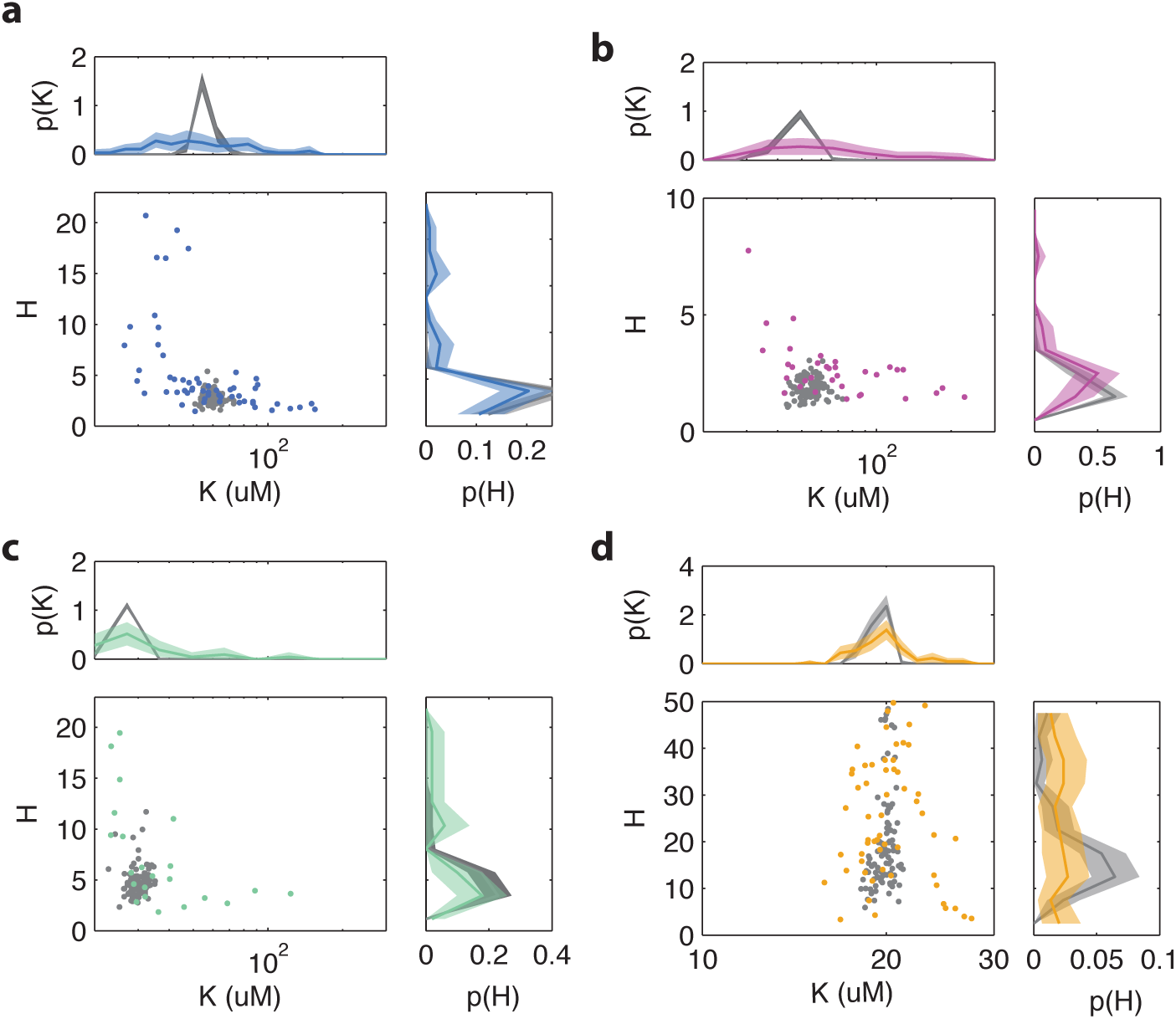
Influence of experimental noise on Fit parameters *K* and *H* from Hill curve Ats to single-cell dose-response experiment. Colored points and lines indicate Fits to measurement data, gray lines and points are from a simulation in which gaussian white noise is added to a dose response curve with *K* and *H* obtained from a Fit to the population averaged time series. The noise level of the simulation is chosen such to approximate the average mean-squared error [MSE] of the dose response curve Fits. Experimental data with a MSE exceeding a determined threshold are removed from the analysis. For the experiment on cells with WT receptor complement (TSS58) a maximum mse of 0.05 is used, excluding 5 cells for (**b**) for the experiment at OD=0.45, this excludes 5 cells, at (**d**) OD=0.31, 3 cells and at (**c**) OD=0.59, 6 cells. (**d**) For the experiment on Tsr-WT (UU2567) 11 cells are removed by the same criterion. All shaded areas indicate 95 % confidence intervals obtained through bootstrapping.

**Figure 3 - Supplement 1.**
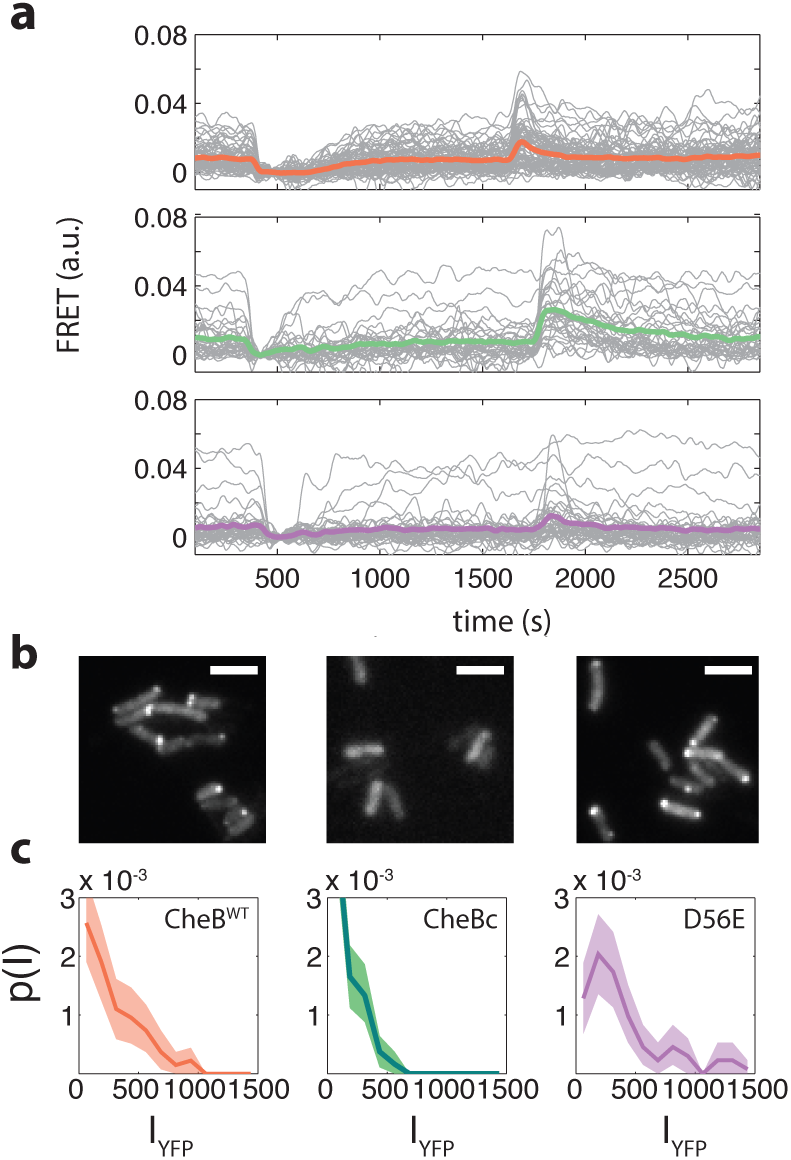
**a** Example time FRET time series for cells expressing CheB^WT^ (top, pink), CheB^D56E^ (bottom,purple), CheB_c_ (middle, green) in ΔCheB (VS124) background. **b** Localization of CheB in the cell probed by mVenus fusions to CheB. Clustering was quantified by the fraction of cells clearly showing one or more clusters in the cell. Shown are representative examples of CheB^WT^ (left, 78% clustered), CheB_c_ (middle, 0%), CheB^D56E^ (right, 72%). Scale bars 2 μM. **c** Histograms fluorescence intensity for single-cells in units of photons/pixel. Mean and standard deviation of these distributions for the strains is (from left to right) 272 ± 237 (114 cells), 148 ± 133 (150 cells) and 398 ± 328 (106 cells). This corresponds to *CV* 0.87, 0.90, 0.82 for the different CheB genotypes.

**Figure 3 - Supplement 2.**
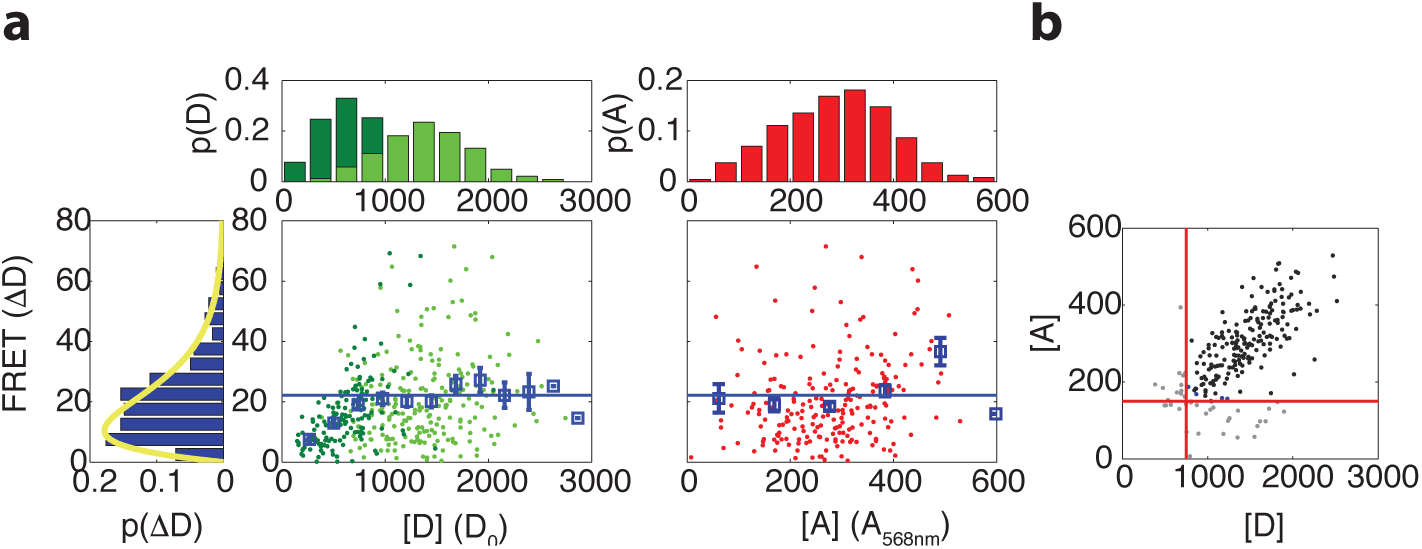
Relation between maximum FRET response and FRET fluorophore expression levels. a) (Left) Scatter plot with donor intensity (*D*_0_, green) versus FRET response to 500 *μM* MeAsp (ΔD) and (right) acceptor intensity (A, red, measured by direct excitation) with marginal distributions for *A, D* and *ΔD* in VS104/pSJAB12. For the donor intensity we measured at low induction (10 μΜ IPTG, dark green) and high induction (100 μΜ IPTG, light green). The acceptor intensities were only measured at high induction. In the scatter plots the mean FRET response is plotted with the error bars (blue), the horizontal line indicating the average for the binned data as described in panel (**b**). The yellow curve is a fit to a gamma distribution with corresponding values and 95% confidence intervals of *k* = 2.0 ± 0.4 and *Θ* = 10.2 ± 2.5. All fluorescent intensities are measured in photons/pixel. All histograms are normalized to the number of cells. b) Gating of the data. Scatter plot of donor intensity versus acceptor intensity. All data with fluorescent intensities (in photons/pixel) lower than D=750 or A=175 are excluded from the analysis since the FRET response depends on donor and acceptor intensity levels below these levels. The distribution of the FRET response (extreme left) is only for the gated data.

**Figure 3 - Supplement 3.**
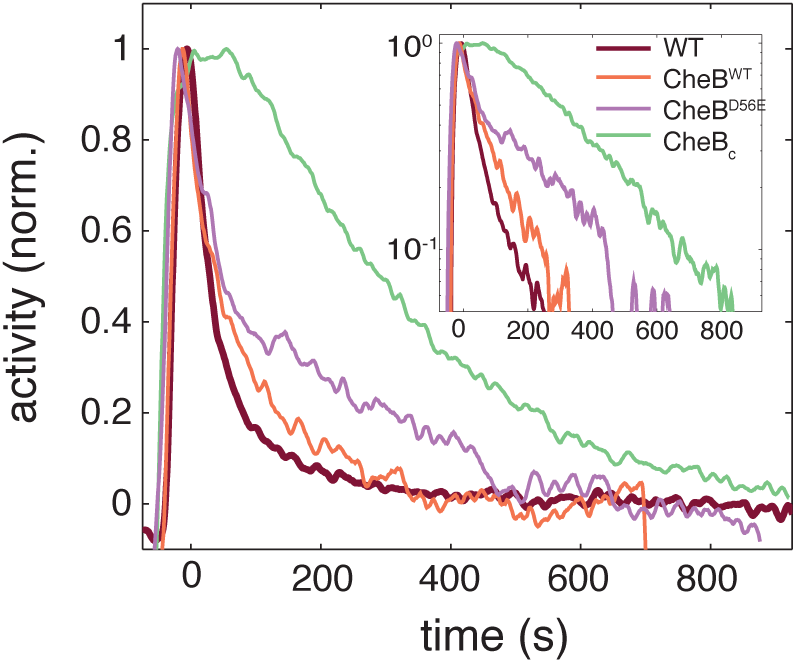
Phosphorylation feedback is not a necessary condition for fast removal adaptation dynamics. Population FRET time series of cells expressing CheB^WT^ (orange), CheB^D56E^ (purple), CheB_c_ (green) in ΔCheB (VS124) background as well as WT (CheB from native chromosome position, VS104, brown) are shown after removal of 500 μM to which cells have adapted. The strain expressing CheB^D56E^ lacks phosphorylation feedback but has a fast removal response. The population FRET experiment is performed as described previously (***Sourjik and Berg, 2002b***). The strains and induction levels are the same as in the single-cell FRET experiments on the CheB mutants.

**Figure 3 - Supplement 4.**
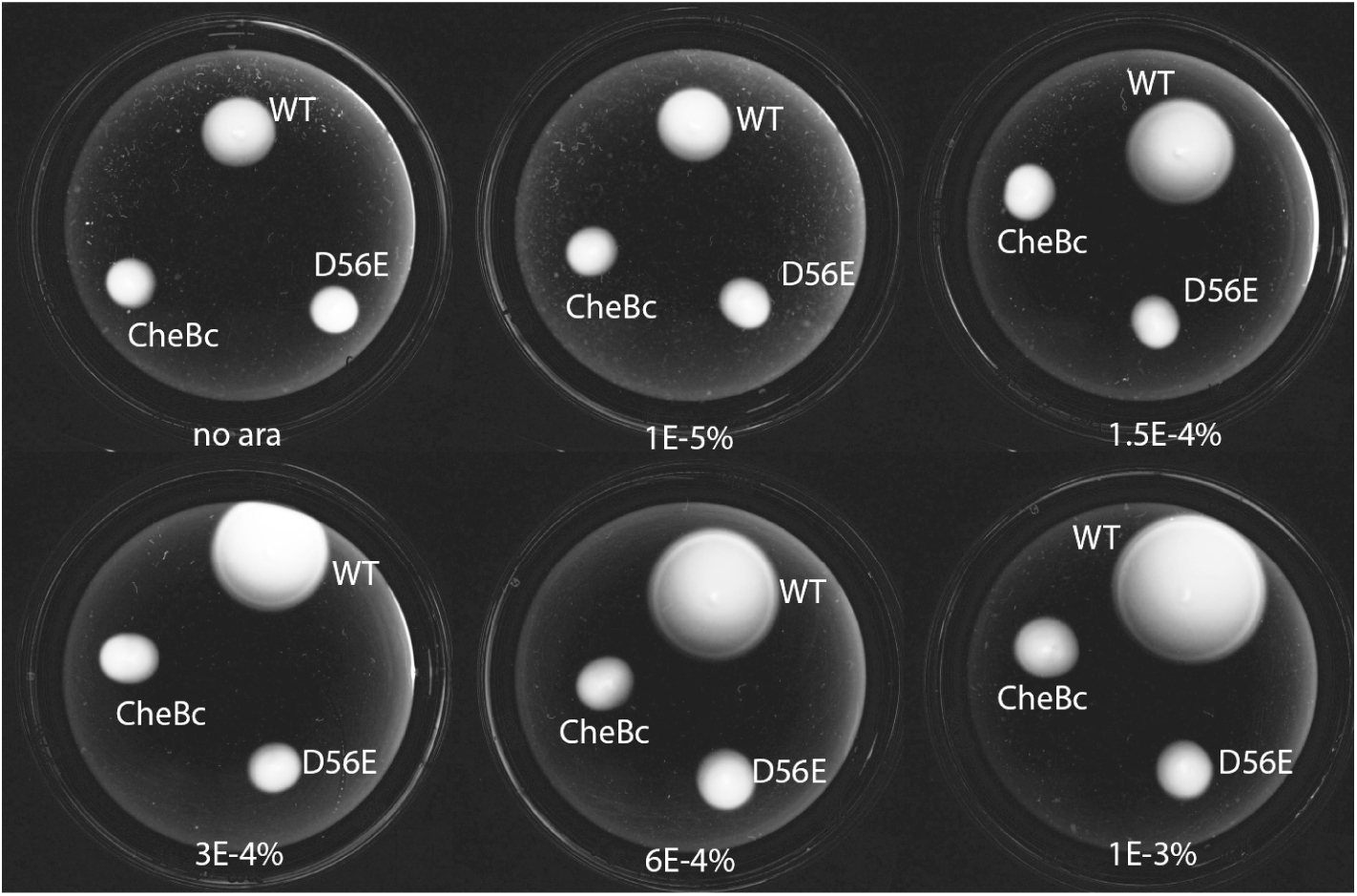
Phosphorylation defective mutants show impaired chemotaxis on soft agar. Dark-Field images after 14 h of growth and motility on soft agar plates (0.26 % agar in TB with appropriate antibiotics, kept at 33.5*^0^* C). The different strains express either WT CheB, CheB-D56E and CheBc from an arabinose inducable pBAD plasmid in Δ CheB strain (UU2614). The arabinose concentration is varied from 0 % to 0.001 %

**Figure 4 - Supplement 1.**
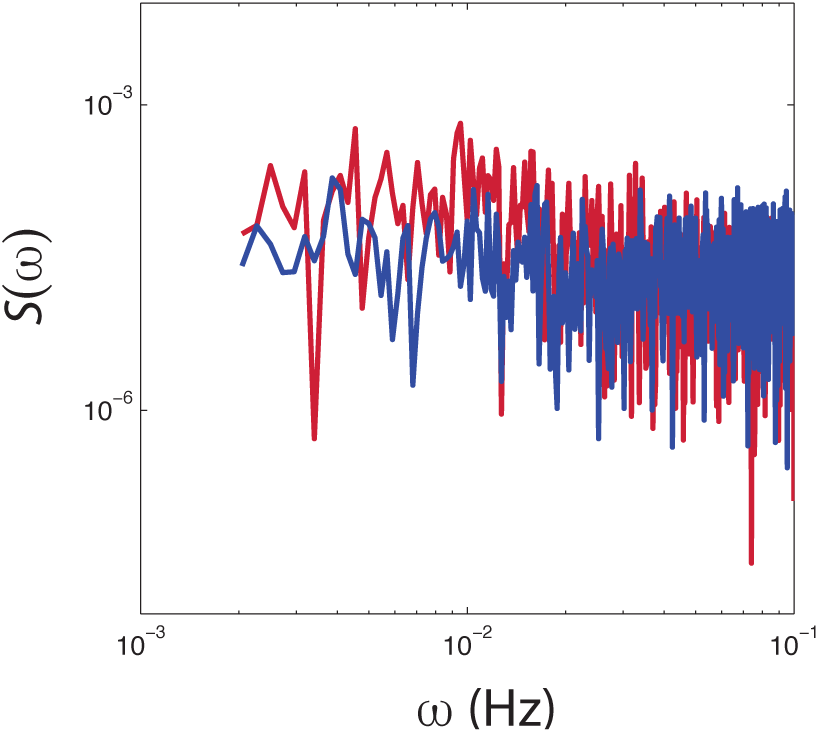
Power spectral density estimates from population averaged time series of CheRB+ (red) and CheRB-cells (blue). The time series are from the same experiment as shown in Figure 3b

**Figure 4 - Supplement 2.**
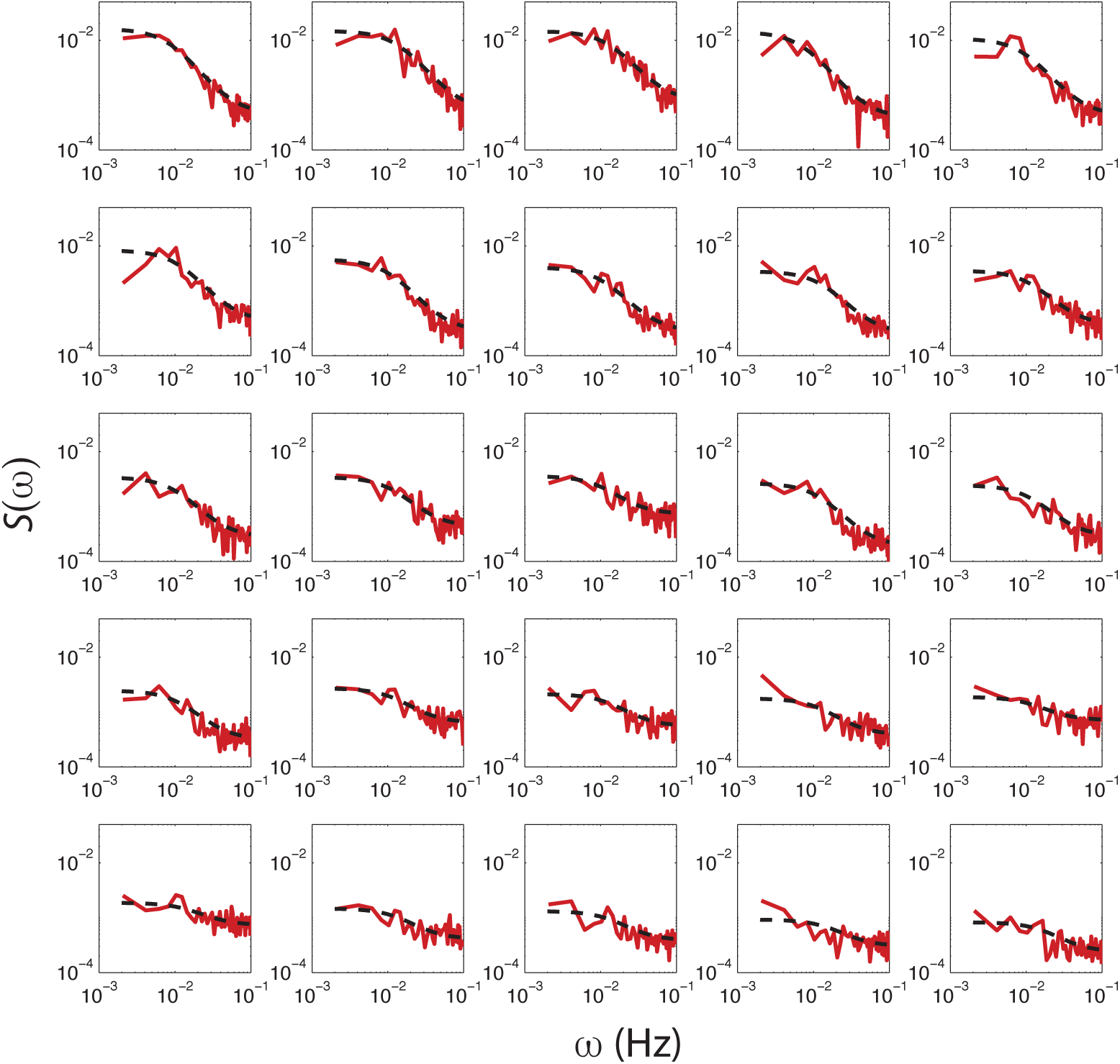
PSD estimates obtained from single-cell FRET time series (red dashed curve) with Fits of O-U process to PSD estimates to 25 out of 31 cells (black dashed curve) from a single experiment shown in Figure 3b. The cells are sorted by variance calculated from the Fit (cτ/2), with top-left having the highest. Cells without a clear low frequency plateau, higher than Five times the standard deviation of the high frequency noise, were excluded from the analysis.

**Figure 4 - Supplement 3.**
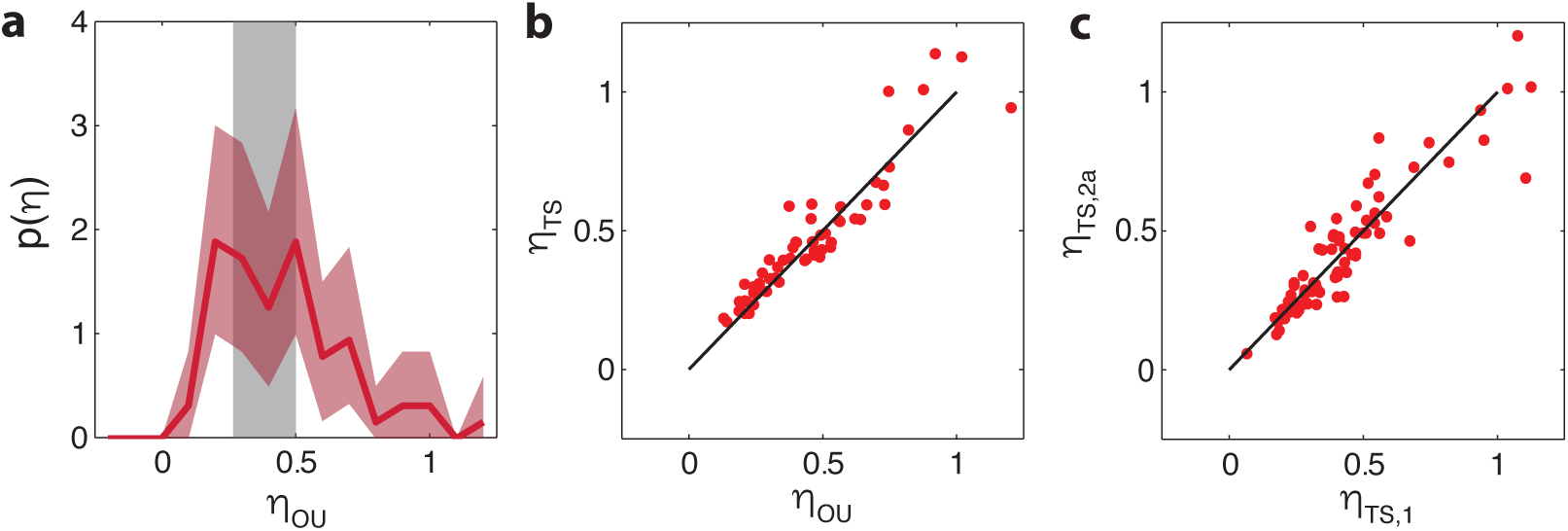
Comparison between noise amplitudes obtained from time series and power spectra. **a** Noise amplitude 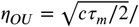 〈*a*〉 for 75 CheRB+ cells time series (from 3 experiments) obtained with mean noise level of 0.42. 14 cells without a clear low frequency noise plateau, at least five times the standard deviation of the high frequency noise floor of the power spectrum, were excluded from the analysis because *τ* could not be constrained by the fit. The grey shaded area indicates the expected noise amplitude variability based on a f nite time window and experimental noise in the experiment (see **Materials and Methods**). The width is def ned as the mean plus (minus) one standard deviation of the distribution of noise amplitudes obtained from simulated OU time series. **b** Noise amplitude *η_OU_* obtained from fits of Ornstein-Uhlenbeck process to power spectra versus the noise amplitude *η_TS_* obtained through calculating the standard deviation of the FRET time series after 10s average filtering. Also shown is the diagonal *η_OU_* = *η*_TS_. **c** The noise amplitude of the f rst (*η_TS_*,*1*) and second half (*η_TS_*,*_2_*) of the time series show high correlation, indicating that fluctuations are constant throughout the experiment. All noise amplitudes are def ned as coefficient of variance.

**Figure 5 - Supplement 1.**
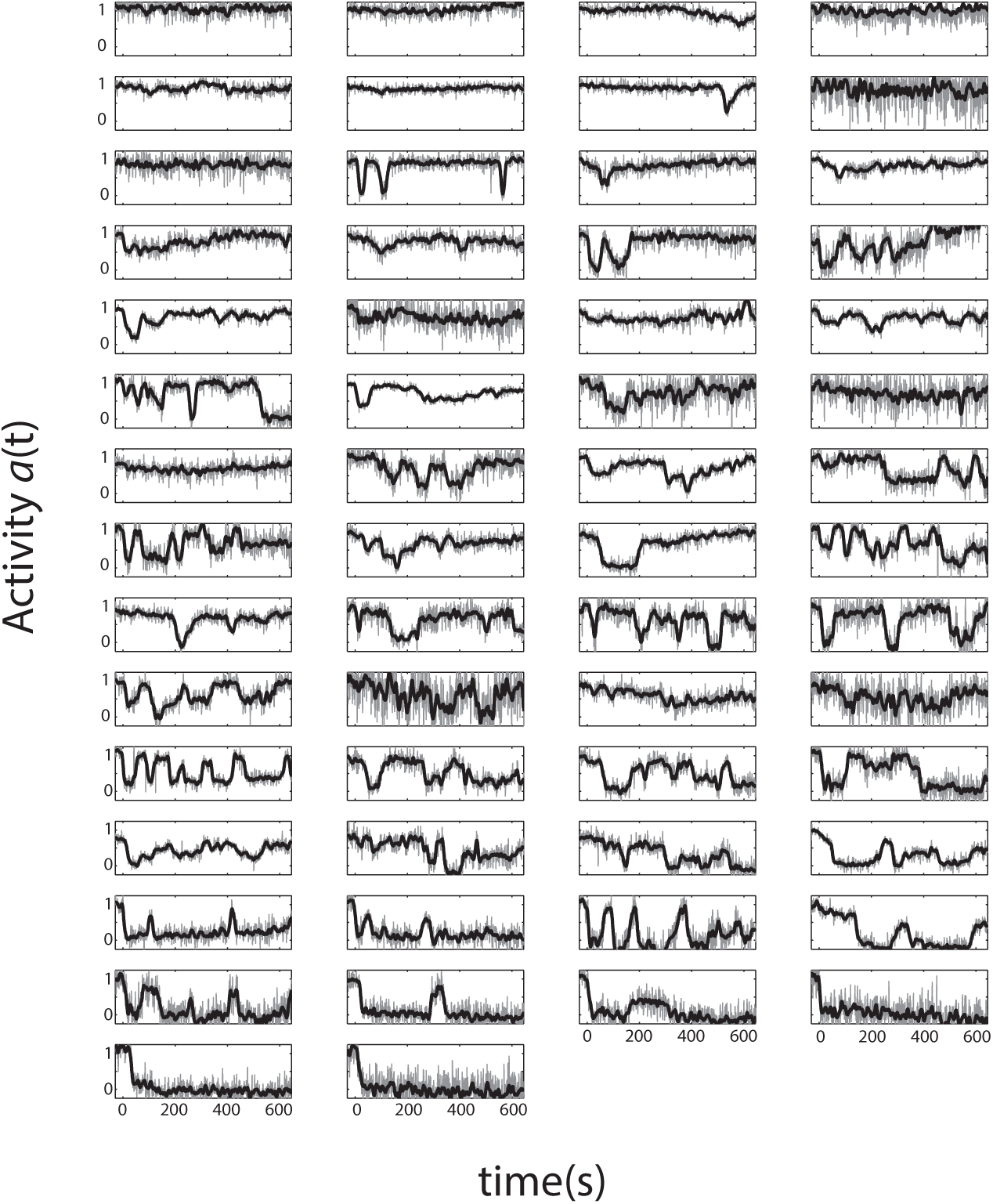
Single-cell activity time series from a step response FRET experiment. Shown are 58 cells with Tsr as the only chemoreceptor in CheRB-background (TSS1964) exposed to a step stimulus of 20 μM L-serine, each normalized to their own response amplitude. Time 0 is set to the time at which the population-averaged signal starts responding to a stimulus. The panels are ordered by the average activity level over the plot range.

**Figure 5 - Supplement 2.**
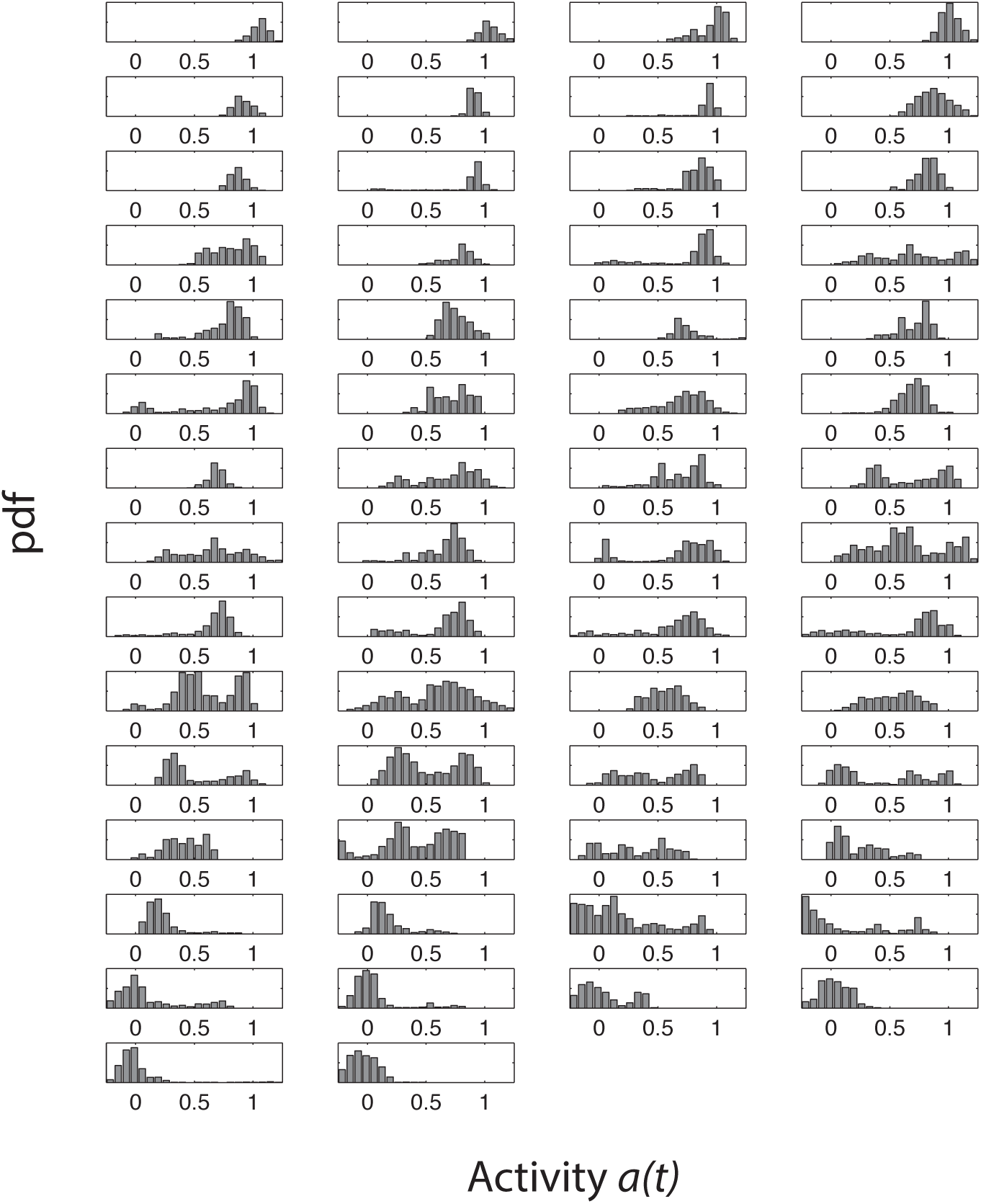
Single-cell activity histograms series from a step response FRET experiment. Shown are 58 cells with Tsr as the only chemoreceptor in CheRB-background (TSS1964) exposed to a step stimulus of 20 μM L-serine. The order of the plots is the same as in Figure 5 - **Supplement 1**

